# Allele pairing at Sun1-enriched domains at the nuclear periphery via *T_1_A_3_* tandem DNA repeats

**DOI:** 10.1101/2023.04.07.536031

**Authors:** Gayan I. Balasooriya, David L. Spector

## Abstract

Spatiotemporal gene regulation is fundamental to the biology of diploid cells. Therefore, effective communication between two alleles and their geometry in the nucleus is important. However, the mechanism that fine-tunes the expression from each of the two alleles of an autosome is enigmatic. By developing an allele-specific gene expression visualization system in living cells, we show that alleles of biallelically expressed *Cth* and *Ttc4* genes are paired prior to acquiring monoallelic expression. We found that active alleles of monoallelic genes are preferentially localized at Sun1-enriched domains at the nuclear periphery. These peripherally localized euchromatic DNA loci harboring monoalleleic genes are enriched with adenine-thymidine-rich tandem repeats, which are bound by Hnrnpd, the “molecular wrench”, to localize these loci in a Hi-C-defined A compartment within the B compartment. Our results demonstrate the biological significance of *T_1_A_3_* tandem repeat sequences in genome organization and how the regulation of gene expression, at the level of individual alleles, relates to their spatial arrangement.

## Introduction

Numerous studies have shown that chromatin is organised into nuclear territories in the interphase mammalian nucleus, seemingly linking chromosome architecture to gene regulation and transcriptional heredity^1–7^. In a diploid genome, precise regulation of transcription dosage requires effective communication between the two alleles; therefore, the positioning of the two alleles of a gene in the nucleus may play a significant role in “alleleic communication” in the regulation of autosomal gene expression^8–13^. Although in theory, both alleles of a gene could be equally exposed to the identical *trans* environment in the nucleus, the presence of nuclear domains indicates a compartmentalized nucleoplasm, potentially facilitating gene-gathering in “hubs” owing to genome folding and allele pairing potential^14–18^, which could significantly impact the regulation of allelic-specific gene expression. Autosomal monoallelic gene expression has been identified in mammalian diploid cells in which only a stochastically or deterministically chosen single allele is transcriptionally active in a given cell^19–21^. Although a recent study showed allele-specific differential regulation of monoallelic genes^22^, the mechanism that regulates the allele choice of monoallelic genes, which is distinctive to gene imprinting, and the allelic geometry of those genes remain poorly understood.

Pioneering molecular techniques have been developed to visualize mRNA, DNA, and proteins in live cells, advancing our understanding of the spatial and temporal dynamics of those molecules^23–31^. However, the techniques available to date for exploring genome regulation in living cells can not capture the spatio-temporal dynamics of allelic gene expression, especially referring to the parental origin of transcripts and alleles simultaneously, precluding a deep understanding of the molecular mechanisms of monoallelic gene expression regulation in a context-dependent manner. Therefore, developing molecular tools and techniques to gain mechanistic insights into spatiotemporal allelic gene expression regulation in living cells will immensely deepen our fundamental understanding of the governing principles of gene expression in organismal development, homeostasis and disease^32–34^.

In this study, we developed a molecular tool to distinguish the spatiotemporal expression of RNAs, referring to their maternal and paternal alleles’ spatial organization in a living cell nucleus, using polymorphic F1 hybrid mouse embryonic stem cells (mESCs). Utilizing this system to probe two monoallelically expressed autosomal genes in neural progenitor cells (NPCs), we showed that when the genes are biallelically expressed in mESCs, alleles are paired, and the transcriptionally active alleles are localized at Sun1-enriched domains at the nuclear periphery (NP). We further showed, upon mESCs differentiation to NPCs, that those genes gain monoallelic status, and the active alleles remain localized at the Sun1-enriched domains. By immunoprecipitating the chromatin associated with Sun1-enriched domains, we further showed that monoallelic genes are preferentially expressed at the Sun1-enriched domains at the NP compared to biallelic genes. Using publicly available data, we further showed that the monoallelic genes are localized in “micro-TADs” in Hi-C-defined euchromatic A compartments within peripherally located LADs. In addition, we discovered that genes localized at the NP are associated with *de novo T_1_A_3_* tandem DNA repeat sequences and that these *T_1_A_3_* repeats are involved in monoallelic gene localization to the Sun1-enriched domains by binding to Hnrnpd.

## Results

### Visualizing allele-specific gene expression in living cells

Live imaging of subnuclear structures is challenged by molecular noise inherent to biological systems^31,35,36^, which significantly impacts image processing and analysis due to the low signal-to-noise ratio. In addition, ectopically expressed fluorescently labelled reporter proteins may cause molecular crowding in cells, potentially altering cellular physiology. Furthermore, cell-penetrating fluorescent reporter molecules^37^ may, in some cases, become a limiting factor for long-term time-lapse imaging. To overcome these limitations, we first generated a doxycycline-inducible, tunable EYFP gene fused to the MS2 coat protein (MS2-EYFP) and an mCherry gene fused to the PP7 coat protein (PP7-mCherry) expressing mESC line (Fig. 1a and Extended Data Fig. 1a) using a CRISPR-based approach. Second, we conditionally inserted 24 PP7 stem-loop repeats (PP7-SL) sequence into the 3’ UTR of the CAST/EiJ allele (paternal) and 24 MS2 stem-loop repeats (MS2-SL) sequence into the 3’ UTR of the C57BL/6J allele (maternal) of two different genes using Cas9-D10A nickase and allele-specific SNP-targeting gRNAs (Fig.1a). We established this allele-specific gene expression visualization system for two random autosomal monoallelic (RaMA) genes, *Cth* and *Ttc4*, which are biallelically expressed in mESCs but become monoallelic upon differentiation^22^, using two independent mESC clones (Fig. 1c).

**Figure 1.**
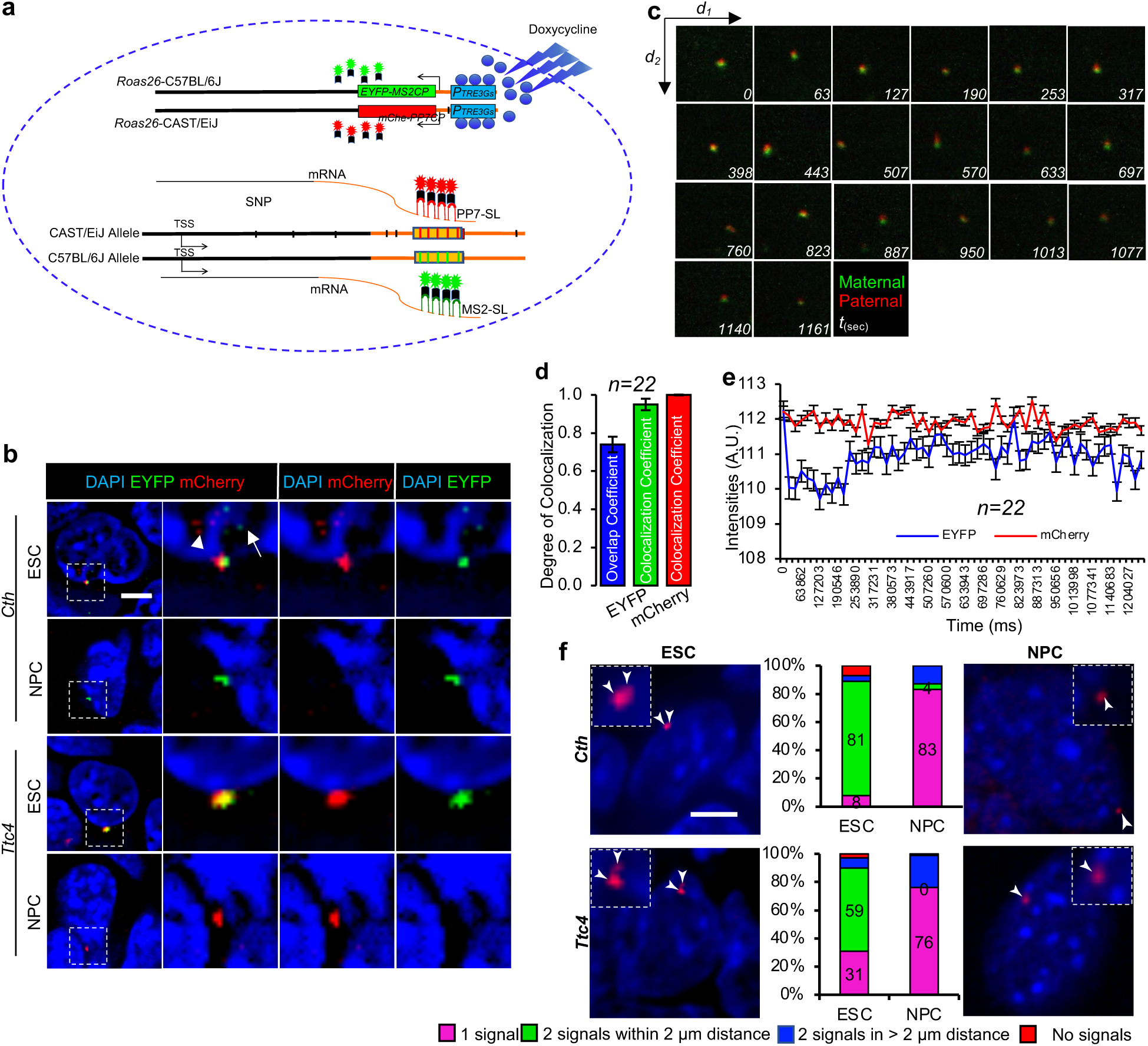
Development of an allele-specific gene expression system to assess the localization of *Cth* and *Ttc4* alleles. **a**, Schematic illustration of the allele-specific gene expression visualization system. Inducible expression of EYFP-MS2CP and mChe-PP7CP is sensitive to doxycycline dosage. These coat proteins interact with unique stem-loops, MS2-SL with EYFP-MS2CP and PP7-SL with mChe-PP7CP at the 3’ UTR of nascent mRNAs. Each RNA molecule contains 24 stem-loops. The maternal (C57BL/6J) mRNAs are marked by EYFP, and the paternal (CAST/EiJ) mRNAs are marked by mCherry. Low-background fluorescent signals are an advantage in this system, as they enable easier allele detection and do not require significant background noise reduction during image analysis. Additionally, this introduces minimal disturbance to the cell’s physiology because the crowding of fluorescent molecules within the cells is minimal and controllable. **b,** EYFP-MS2CP and mChe-PP7CP signals in doxycycline- induced *Cth* and *Ttc4* ESC and NPC cells. Cells were fixed 36 hours post- doxycycline induction. Dotted areas in the left image panels are enlarged to demonstrate the fluorescent signal colocalisation at the nuclear periphery. Fluorescent signals, possibly from single RNA molecules (maternal: arrow, paternal: arrowhead), are visible in the enlarged images in the first row. Scale bar - 6 µm. **c,** Snapshots of a tracked single pair of alleles at different time points from a 20-minute video using the 3i lattice lightsheet microscope. Green and red dots represent the maternal and paternal *Cth* alleles, respectively. **d,** Degree of colocalisation of the EYFP and mCherry signals calculated from tracked 22 EYFP and mCherry signal pairs in 22 cells over a 20-minute video. Columns represent the overlap coefficient of green and red dots (blue column) and colocalisation coefficients of EYFP (green column) and mCherry (red column). Error bars: SEM. n = 22 cells. **e,** Line graph showing fluctuations in the intensity of the colocalised EYFP and mCherry dots. Live images were captured every minute using the 3i lattice lightsheet microscope. Error bars: SEM. n = 22. **f,** Representative RNA-FISH images indicate localisation of nascent mRNAs of *Cth* and *Ttc 4* alleles at the nuclear periphery in mESCs and NPCs. Arrowheads indicate the nascent RNA-FISH signals. Magnified images (within the dotted-lined box) show active pairing of the *Cth* and *Ttc4* alleles. Scale bar - 6 µm. The column graphs show the percentage of cells quantified for distances between red and green dots in nascent RNA-FISH images. Two alleles within a 2 µm proximity are considered as paired alleles.

**Extended Data Figure 1.**
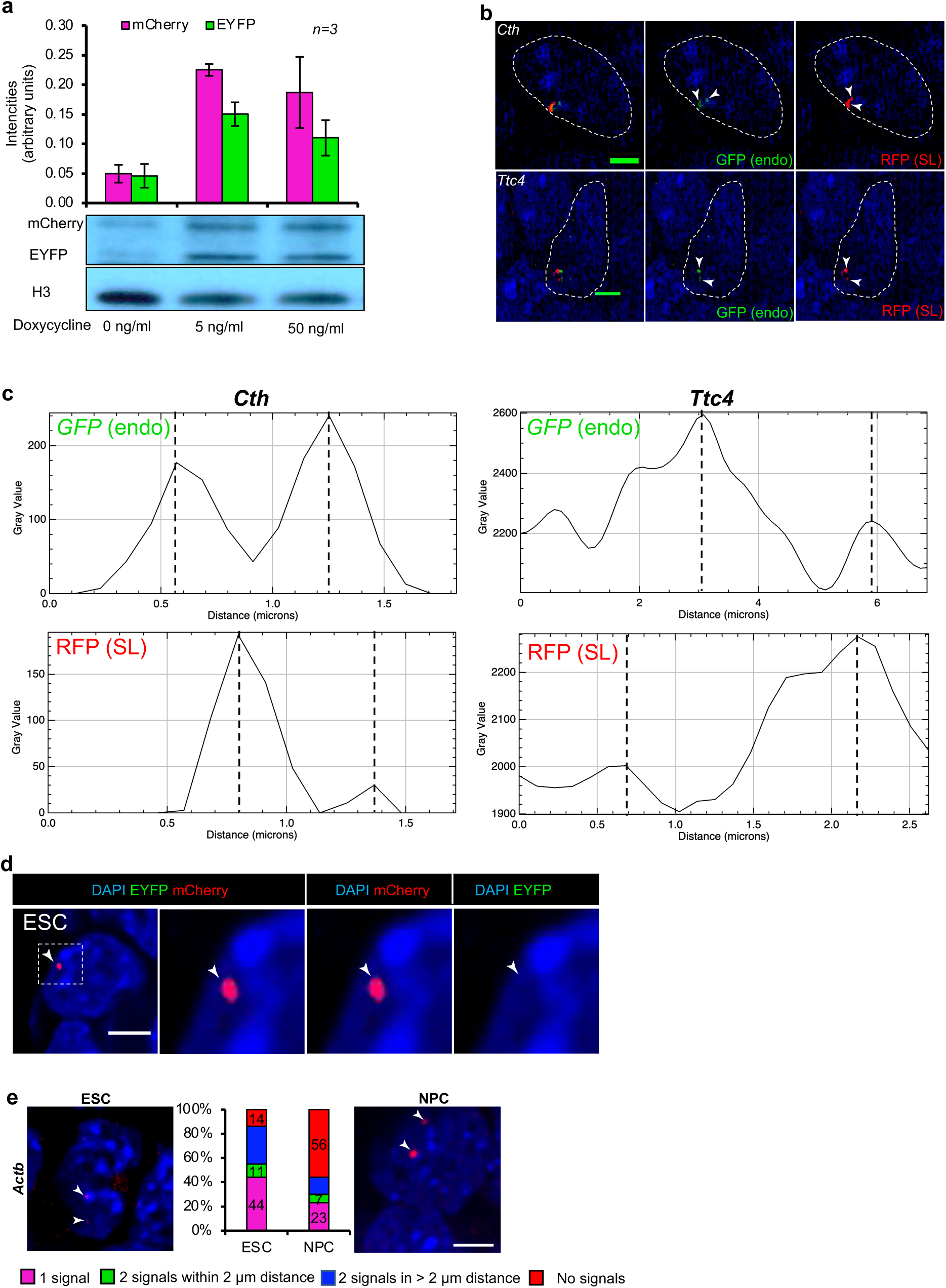
Development of an allele-specific gene expression system to assess the localization of *Cth* and *Ttc4* alleles. **a**, Immunoblot (lower panel) showing EYFP and mCherry responsiveness to doxycycline dosage. Bar graph presenting quantification of the EYFP (using GFP antibody) and mCherry bands in immunoblots from three independent experiments (n=3). **b**, DNA-FISH of *Cth* and *Ttc4* genes targeted with MS2 and PP7 stem-loops in NPCs. Endogenous alleles are labelled with the nick-translated green fluorescent probes, and the stem-loops are labelled with the nick-translated red fluorescent probes. Total DNA is labeled with DAPI. Two signals were detected for green and red in these positively labeled (71%) nuclei. No nuclei exhibited only green or only red signals. Scale bar – 6 µm. **c**, The distance between the two *Cth* alleles and the two *Ttc4* alleles in NPC nuclei. The average distances between the endogenous *Cth* alleles are approximately 0.6 µm, and the two inserted stem-loops are around 0.55 µm, whereas the average distance between the endogenous Ttc4 alleles is about 2.5 µm and the stem-loops approximately 1.5 µm. **d**, Monoallelic expression of the *Ttc4* gene in mESCs. Although *Ttc4* was expected to be expressed biallelically in mESCs, we observed that *Ttc4* is monoallelically expressed in 31% of the mESCs. **e**, Nascent mRNA FISH assay for *Actb* in wild-type mESCs and NPCs. Arrowheads indicate nascent RNA-FISH signals. The accompanying column graphs show the percentages of observations for the distance between the two alleles. Alleles within 2 µm of each other are considered paired. Scale bar - 6 µm. Active *Actb* alleles were paired in only 11% of mESCs, while the remaining cells positive for RNA-FISH signals showed single allelic (44%) and biallelic (11%) expression. Compared to *Cth* and *Ttc4* (Fig.1f), the percentage of cells lacking signals (ESCs: 14% and NPCs: 56%) and the previous observation^22^ of the imbalance of mature versus nascent RNAs of Actb, along with published data^85^, suggest that single allelic expression (44%) of Actb may be due to gene pulsing.

### Active *Cth* and *Ttc4* alleles are paired

To confirm that the MS2-SL and PP7-SL are integrated into the targeted *Cth* and *Ttc4* gene alleles, but not off-targeted to different loci, we performed dual-color DNA-FISH using nick-translated probes designed to detect the endogenous alleles (green) and the inserted stem-loop cassettes (red).

In the IF images, we observed the red and green fluorescent signals exclusively adjacent to each other, confirming on-target stem-loop knock-in for both the *Cth* and *Ttc4* genes (Extended Data Fig.1b). We observed that both alleles of the *Cth* gene were within ∼1µm distance of each other, and the alleles of the *Ttc4* gene were within less than ∼3µm distance of each other in mESCs (Extended Data Fig.1b, c). To assess the coat-protein binding specificity toward their cognate stem-loops, we induced the expression of MS2-EYFP and PP7-mCherry coat proteins in mESCs and mESC-derived NPCs to label nascent RNAs from the *Cth* and *Ttc4* alleles. Imaging of fixed mESCs and NPCs, confirmed stem-loop-coat protein specificity in our knock-in clones (Fig. 1b and Extended Data Fig. 1d).

Homologous allele pairing has been shown to influence gene regulation^8,11,38^, yet the broader impact of allele pairing and three-dimensional nuclear arrangement on transcriptional heredity remains poorly understood. By examining nascent RNAs from the maternal and paternal alleles of the *Cth* gene every 20 seconds over 20 minutes by live-cell imaging (Fig.1c and Supplementary Movie 1), we observed that the two actively transcribing alleles frequently colocalize in mESCs (Fig.1d, e; n=22 cells).

Because genome editing at specific loci can potentially perturb chromatin organization^39,40^, we asked whether the observed allele localization is an artifect caused by insertion of stem-loops to both alleles. To test this, we performed nascent RNA-FISH for the *Cth* and *Ttc4* genes in wild-type mESCs and mESC-derived NPCs (Fig.1f and Extended Data Fig.1e). By quantifying the distance between flurosent signals (nasent RNA-FISH), we observed that the *Cth* alleles are paired in 81% of the mESCs (8% are monoallelic), and the *Ttc4* alleles are paired in 59% of the mESCs (Fig.1f - column graph). These results confirm that insertion of MS2 and PP7 stem-loops did not alter the chromatin positioning; therefore, allele colocalizion of the *Cth* and *Ttc4* genes represents a true biological phenomenon.

### Active *Cth* and *Ttc4* alleles localize to the nuclear periphery

In the fluorescent images obtained from the mESC and NPC (Fig. 1b, f) allele-specific gene expression systems, and the nasent RNA-FISH images for the *Cth* and *Ttc4* genes (Fig. 1f), we observed that transcriptionally active *Cth* and *Ttc4* alleles localized to the NP (Fig.2a and Extended Data Fig.2a). Therfore, since *Cth* and *Ttc4* are monoallelic genes, to assess whether the active monoallelic gene alleles localized at the NP is a unique characteristic of this gene class, we quantitatively assessed the nuclear positioning of the transcriptionally active alleles of a panel of 13 RaMA and 14 deterministic monoallelic (DeMA) genes by performing a nasent RNA-FISH assay. We found that the DeMA and RaMA genes we assessed were preferentially localized (DeMA genes 95% on average; RaMA genes 93% on average; n=100 cells per gene) at the NP of NPCs (Fig.2b and Extended Data Fig.2b,c). In line with our observation, previous studies have reported that transcriptionally active olfactory receptor genes, -an autosomal monoallelic gene family-, are also transcribed at the NP^41–43^, suggesting the existance of transcriptionally active monoallelic gene hubs at the NP.

### Monoallelic genes localize at Sun1 territories

Following our observation of the localization of transcriptionally active *Cth* and *Ttc4* alleles at the NP, we asked whether these active alleles associate with specific proteins or protein territories at the inner nuclear membrane that might provide a mechanism for harboring alleles. Sun1, an inner-nuclear-membrane protein (Extended Data Fig.2d), is involved in positioning the nucleus in the cytoplasm by interacting with actin filaments via nesprin^44,45^. In addition, Sun1 is associated with the nuclear pore complex, which plays a role in mRNA transport^46,47^ and in harboring active genes^14,48^. Since Sun1-enriched nuclear domains are associated with stabilizing chromatin architecture^49^, their specific molecular function in gene regulation is unknown. Although overall, the lamina-associated domains at the NP harbor heterochromatin^50^, the transcriptional status of genes specifically associated with Sun1-enriched domains is unknown. Using an immunofluorescent assay for Sun1 protein in *Cth* and *Ttc4* mESC generated clones (Fig.1a), we identified that nascent RNAs from both alleles of the *Cth* and *Ttc4* genes colocalized with Sun1 protein at the NP (Fig.2c and Extended Data Fig.2e). Based on this observation, we hypothesized that monoallelic genes might preferentially localize to Sun1-enriched territories at the NP. To test our hypothesis, we performed Sun1 Chromatin ImmunoPrecipitation (ChIP) in mESCs and *in vitro*-derived NPCs. To validate the specificity of the Sun1 antibody, we performed mass spectrometric (MS) analysis of proteins isolated from the Sun1 ChIP assay. In both mESC and NPC MS datasets, Sun1 was the most significantly enriched protein in the pull-down (Fig. 2d and Extended Data Table 1), confirming the antibody specificity for Sun1, which was subsequently confirmed by immunoblot analysis (Fig.2e).

**Figure 2.**
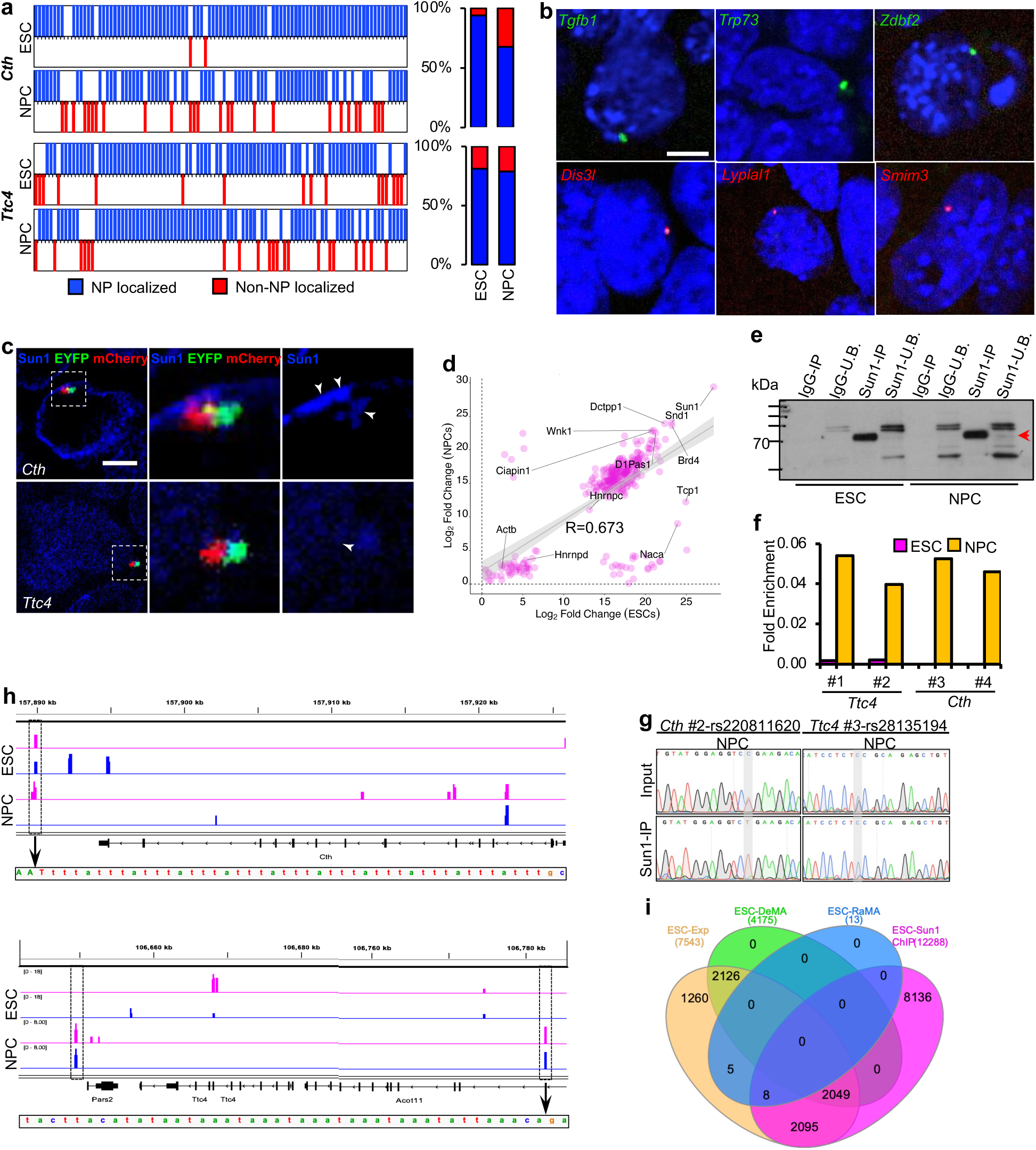
Active alleles are localized at Sun1 protein territories. Quantification of *Cth* and *Ttc4* nascent RNA- FISH signal localization in the nucleus. Each line represents a counted cell, and empty points on the X- axis indicate cells with one signal at the nuclear periphery and the other more internally located in the nucleoplasm. Blue and red lines denote cells with nuclear periphery or nucleoplasm- localized nascent RNA- FISH signals, respectively (n = 100 cells per group). Column graphs display the percentages of RNA-FISH signals at the nuclear periphery versus nucleoplasm in ESCs and NPCs. **b**, Representative nascent RNA-FISH images of transcriptionally active random autosomal and deterministic monoallelic genes we tested in NPCs. Green fonts - imprinted genes, red fonts - stochastic monoallelic genes. Scale bars - 6 µm. **c**, *Cth,* and *Ttc4* genes’ nascent RNA localization at Sun1 protein territories at the NP. Scale bars - 6 µm. **d**, Correlation of proteins pulled down by Sun1 antibody in mESCs and NPCs. The highest enrichment was observed for the Sun1 protein, confirming the antibody’s specificity. The overall Pearson’s correlation coefficient is 0.673, with a 95% confidence interval from 0.606 to 0.73 and a p-value of 3.9e-41. **e**, Immunoblot of the Sun1 pull-down. The Sun1 antibody recognises five Sun1 protein isoforms (lane: Sun1-U.B). **f**, Enrichment of *Cth* and *Ttc4* genes in Sun 1 chromatin pull-downs. Enrichment was assessed via qRT-PCR using primer sets (*Cth*: #1 and #2, *Ttc 4*: #3 and #4 – sequences are in the method section) covering known SNPs. Fold enrichment was calculated relative to input. **g**, Histogram plot of Sanger-sequenced qRT-PCR products from Sun1 chromatin pull-down DNA in NPCs. Parental backgrounds of the alleles were confirmed by SNPs covering primers (*Cth*: rs 220811620- primer set #2 and *Ttc4*: rs 28135194- primer set #3) unique to each allele. **h**, Allele-specific genome browser tracks illustrating Sun1 ChIP-seq peaks. TA-bridge motif-containing peaks associated with the *Cth* and *Ttc4* genes are shown in dotted boxes. Pink lines: Maternal alleles; blue lines: Paternal alleles. **i**, Venn diagram showing the intersections of Sun1-associated genes (including X and Y chromosome genes) identified in ChIP-seq assays and DeMA, RaMA, and all expressed genes in mESCs. RNA expression data are from Balasooriya and Spector, Nat. Commun., 2022^22^.

**Extended Data Figure 2.**
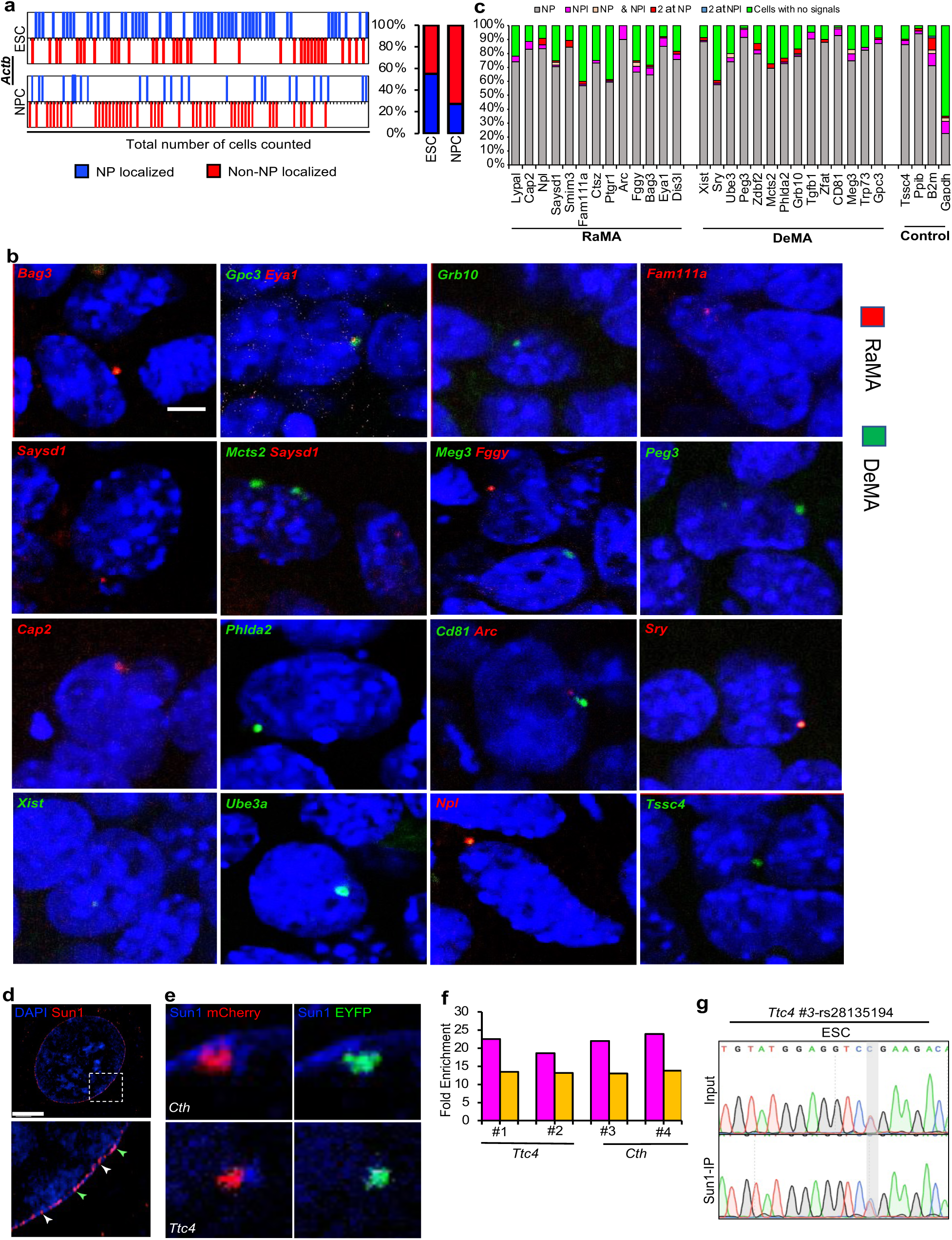
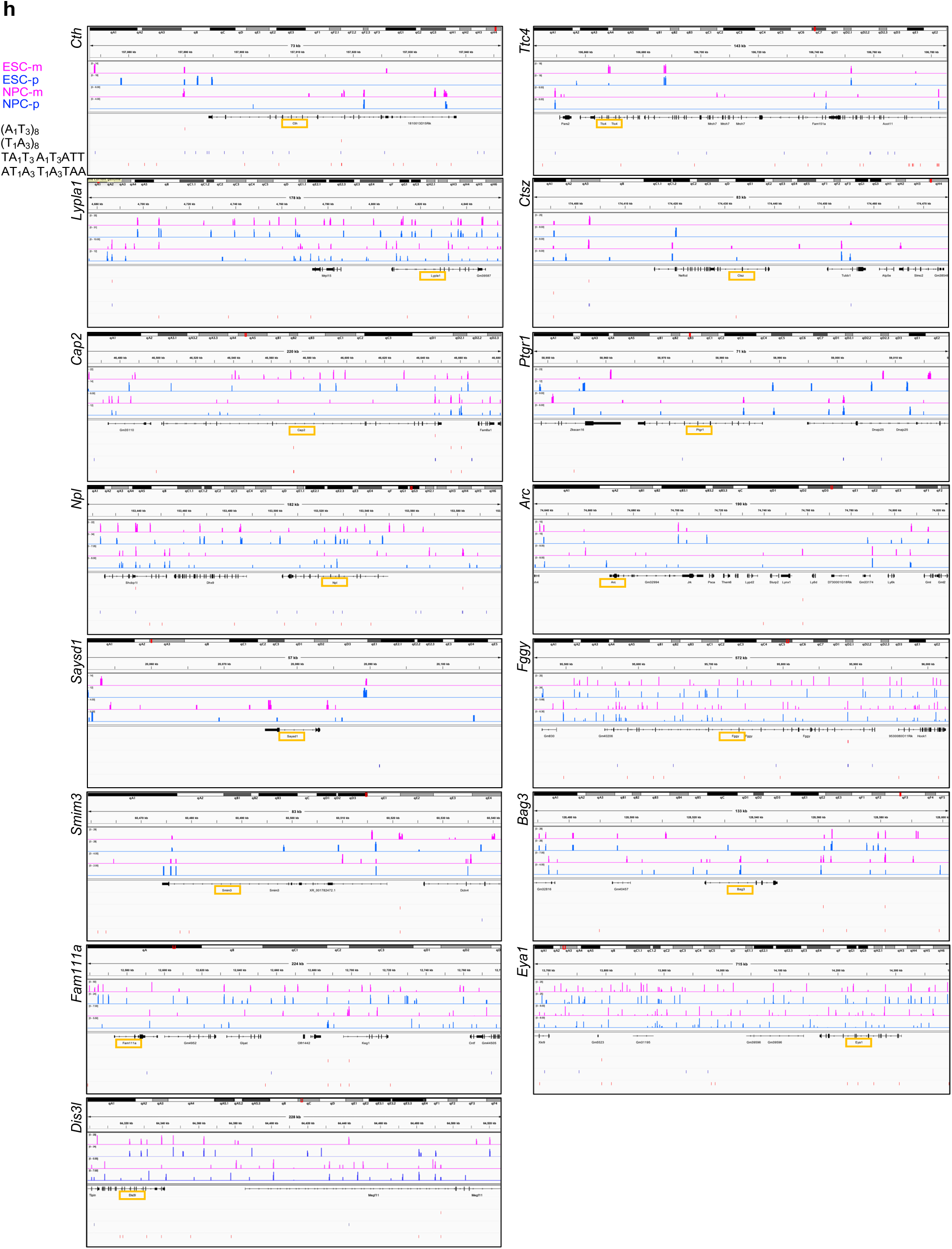
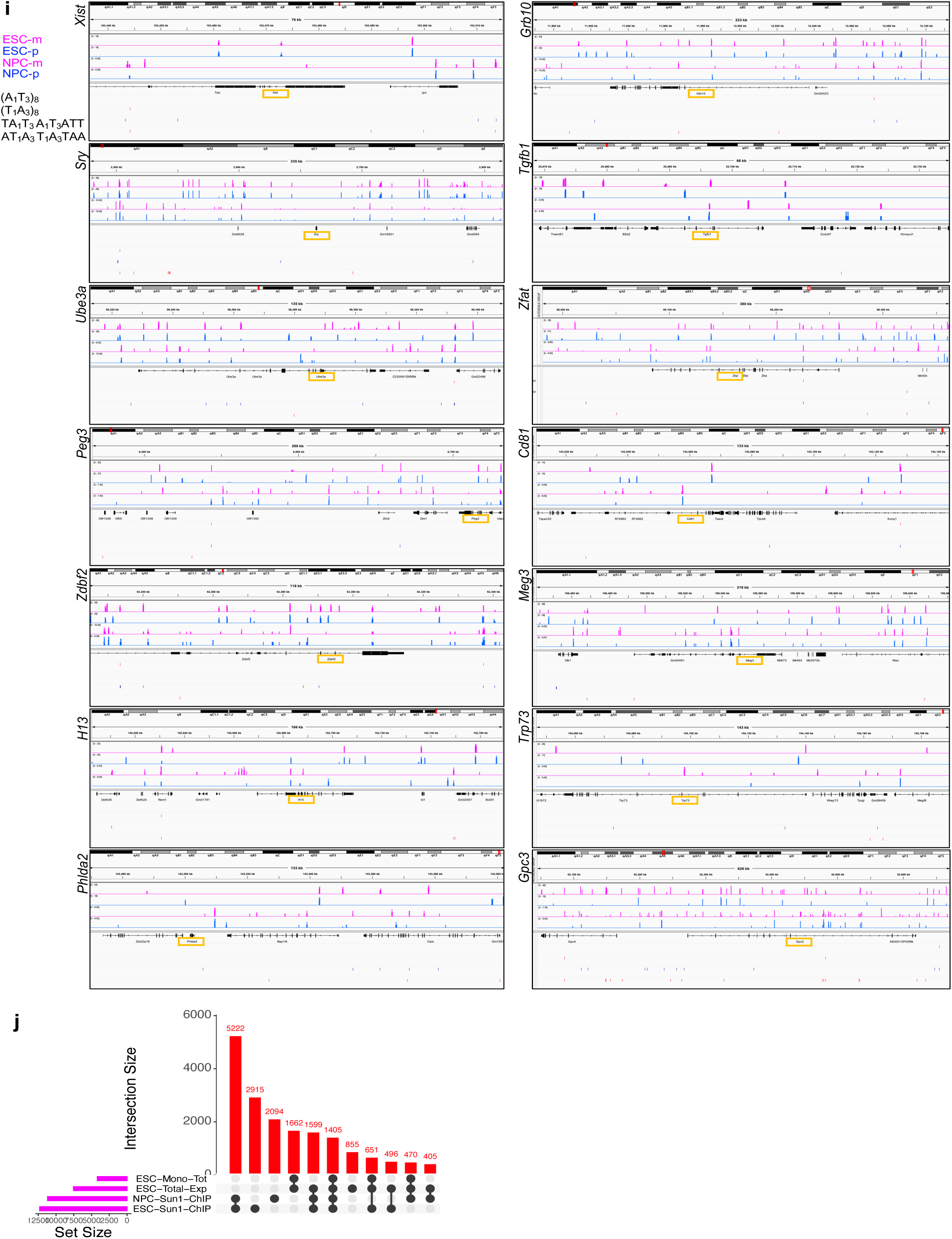
Active alleles of monoallelic and bi-allelic genes are localized at Sun1 protein territories. **a**, Quantification of *Actb* nascent RNA- FISH signals in the nucleus. Each line represents a counted cell, and empty points on the X-axis indicate cells with one signal at the nuclear periphery and the other further inside the nucleoplasm. Blue and red lines indicate nascent RNA-FISH signals localized to the nuclear periphery or nucleoplasm (n=100 cells in each group). **b**, Representative images from the nascent RNA-FISH assay for a panel of monoallelic genes. DeMA genes are labeled in green font, and RaMA genes are labelled in red font. Scale bars - 6 µm. **c**, Quantification of the nascent RNA-FISH signals for RaMA and DeMA genes in NPCs. The Y-axis shows the percentage of cells with RNA-FISH signals located in the nucleus (NP: Nuclear periphery, NPl: Nucleoplasm). **d**, Immunofluorescent image showing Sun1 protein localization at the nuclear periphery. The enlarged dotted area is shown in the lower panel. Green arrowheads indicate Sun 1 and DNA overlapped loci. White arrowheads indicate Sun 1 with no DNA overlapping. Scale bars - 6 µm. **e**, *Cth* nascent RNAs and Sun1 protein, and, *Ttc4* nascent RNA and Sun1 protein localization at the nuclear periphery. Scale bars - 200 nm. **f**, Fold enrichment of Sun1 pull-downed *Ttc4* and *Cth* genes. The same primer sets were used as in Fig. 2f. Fold enrichment was calculated relative to each gene’s DNA fragment enrichment in the input. **g**, Histogram of the Sanger sequenced qRT-PCR products from Sun1 pull-down chromatin from mESCs using *Ttc4* primers (rs28135194). Grey areas indicate the SNPs. Both maternal and paternal alleles were pulled down by Sun1. **h** & **i**, Allele-specific genome browser tracks showing the ChIP-seq peak association of RaMA (**h**) and DeMA (**i**) genes. Pink tracks: maternal alleles; blue tracks: Paternal alleles. The positions of the shortest TA-bridge motifs (n=2) and the longest (n=8) TA-bridge motifs are shown in the panels below the allele tracks. (n: number of motifs). **j**, UpSet plot showing the gene intersections between Sun1-tritories associated genes in ESCs and NPCs, along with the total genes expressed and monoallelic genes (DeMA and RaMA) expressed in ESCs.

By performing qPCR analysis for the DNA isolated from chromatin pulled down by the Sun1 ChIP assay followed by Sanger sequencing, we confirmed that both *Ttc4* alleles are localized to Sun1-enriched territories in mESCs, whereas only a single allele was associated with Sun1 in NPCs (Fig.2f,g and Extended Data Fig.2f, g). Although only a single *Cth* allele was detected in the Sun1 pull-down in NPCs (Fig.2g), the *Cth* gene could not be amplified from the Sun1 pull-down DNA in mESCs (Fig.2f), likely due to low enrichment of *Cth* in the Sun1 pull-down chromatin fraction. Therefore, to assess *Cth* and other genes associated with Sun1-enriched chromatin, we deep-sequenced the DNA fragments isolated from the Sun1-ChIP chromatin. The allele-specific ChIP-seq data showed that the *Cth* peaks were enriched on both alleles in mESCs and on a single allele in NPCs, located at ∼4.5 kb downstream of the *Cth* gene (Fig.2h – upper panel). In NPCs, a single peak was homologously enriched within the intronic region of both *Cth* alleles. For *Ttc4,* allele-specific ChIP-seq peaks were enriched at distinct genomic regions in mESC versus NPCs: in mESCs, peaks localized to the gene body of *Ttc4 or* neighboring gene *Mroh7*, whereas in NPCs, peaks occurred ∼4 kb downstream of the *Fam151a* and *Acot11* gene bodies (Fig.2h – lower panel). In addition, we found that the monoallelic genes we assessed in the nascent RNA-FISH assay were also enriched in the Sun1-associated chromatin (Extended Data Fig. 2h,i). Moreover, we also found that 49% of transcriptionally active DeMA genes and 61.5% of RaMA genes in mESCs (from our previous study^22^) overlapped with the genes enriched in Sun1-associated chromatin (Extended Data Table 2), whereas only 27.8% of transcriptionally active biallelic genes were Sun1-associated (Fig.2i). Furthermore, when we assessed how common these monoallelic gene associations were in Sun1 territories in NPCs, we found that 74.1% of monoallelic genes in mESCs are enriched in the Sun1 chromatin pull-down (Extended Data Fig.2j and Extended Data Table 3).

### “micro-TADs” within LADs harbor active genes

Although lamina-associated domains (LADs) have been shown to harbor heterochromatin^51–54^, emerging evidence suggests that some of the genes within LADs are transcriptionally active^55,56^. Since we found that the active alleles of monoallelic genes were preferentially localized at the NP, we examined whether the Cth, Ttc4, and the monoallelic gene panels we used in the nascent RNA-FISH assay were associated with Hi-C-defined compartments in ESCs and NPCs^57,58^ using the published data in the 4DN data portal^59,60^. We found that, compared to low-resolution compartments, in high-resolution compartments defined by an insulation score-diamond, the Cth and Ttc4 genes (Fig.3a, Extended Data Fig.3a,b) and the monoallelic gene panel we used (Extended Data Fig.3c,d) were housed in “micro A” compartments within B compartments. This finding confirmed our observation of active monoallelic genes at the NP and demonstrated the presence of euchromatic compartments within LADs, which we named “micro-TADs”.

**Figure 3.**
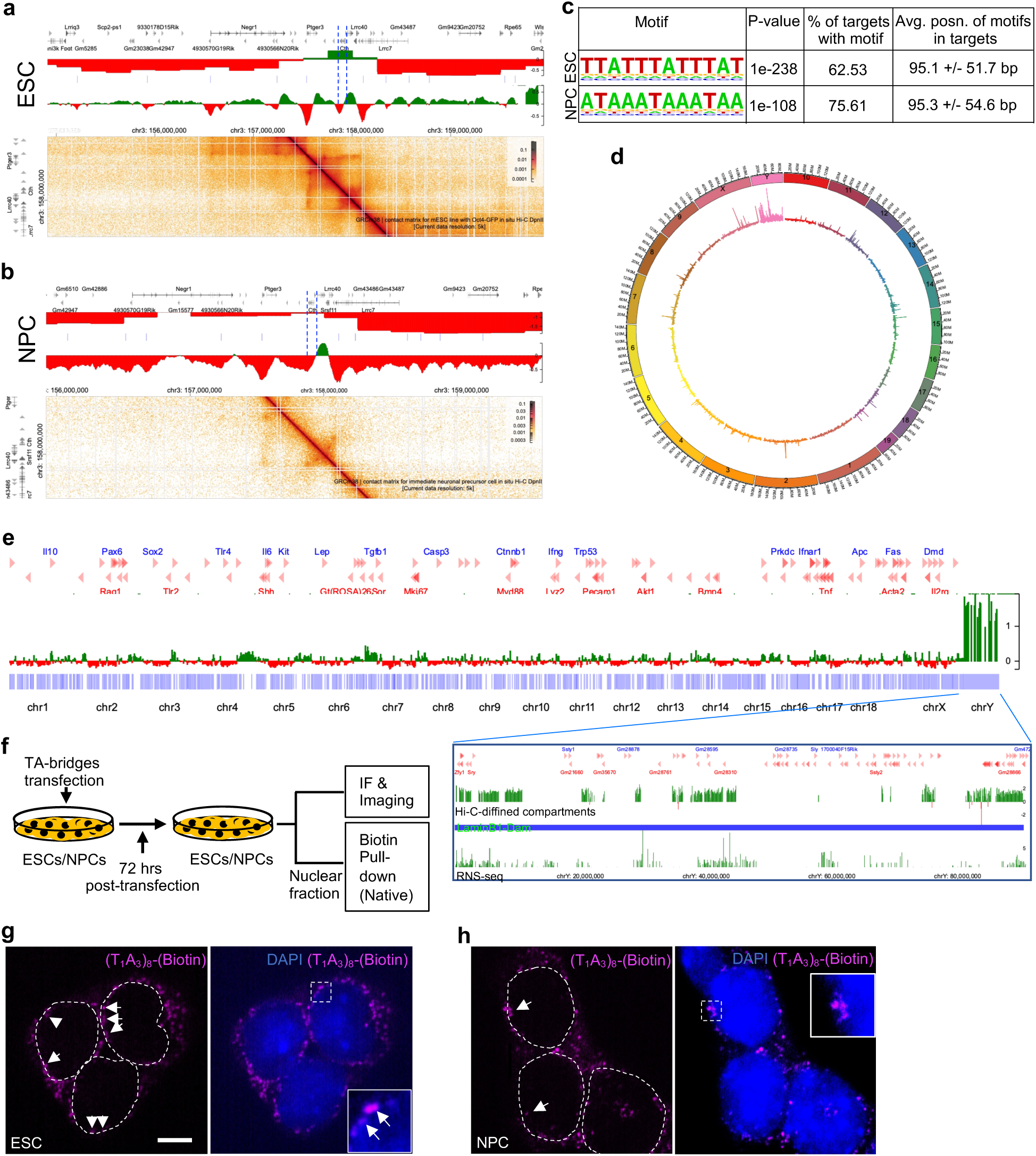
TA-bridges position the *Cth* gene in micro-TADs within the chromatin B compartment. **a** & **b**, *Cth* positioning in Hi-C-defined “micro-TADS”. The *Cth* gene is located in the A compartment (green) within the predominantly represented B compartment (red). The contact Matrix (lower panels) for each gene shows the broader TADs and the “micro-TADs” within the TADs. Vertical lines indicate the boundaries of the TADs. Upon ESCs differentiation into NPCs, the monoallelically expressed *Cth* allele forms an isolated A compartment within the B compartment. Data is from references 59, 60, 61 and 62. **c,** *De novo* TA-bridge DNA motif. **d,** genome-wide association of TA-bridges shown in circular genome representation. The Y chromosome is highly enriched with TA-bridges. The height of the lines represents the motif score at a specific genomic region. Motif analyzis was performed using the Motif Cluster Alignment and Search tool (MCAST) with the following parameters: Sequence length: 11 bases, motif hit threshold: p-value < 5.0E-4, adjacent motif spacing: less than or equal to 50, and match threshold: E-value < 10.0. **e,** Genome-wide Hi-C-defined insulation score-diamond A (green) and B (red) compartment distribution in ESCs. The marked lower panel shows a magnified view of the Y chromosome, including its insulation score-diamond (upper track), LaminB1-DamID (middle track), and RNA-seq read count density (lower track). **f,** Schematic diagram showing the development of the assay used to visualize the TA-bridges spatial localisation and TA-bridge interacting (*in vivo*) proteins. **g** & **h,** Transiently transfected (T_1_A_3_)_8_ single-stranded DNA motif localization in nuclei. mESCs and NPCs were transiently transfected with biotin-tagged (T_1_A_3_)_8_ oligonucleotides. At 72 hrs post-transfection, cells were fixed and immunolabeled for biotin, and DNA was counterstained with DAPI.

**Extended Data Figure 3.**
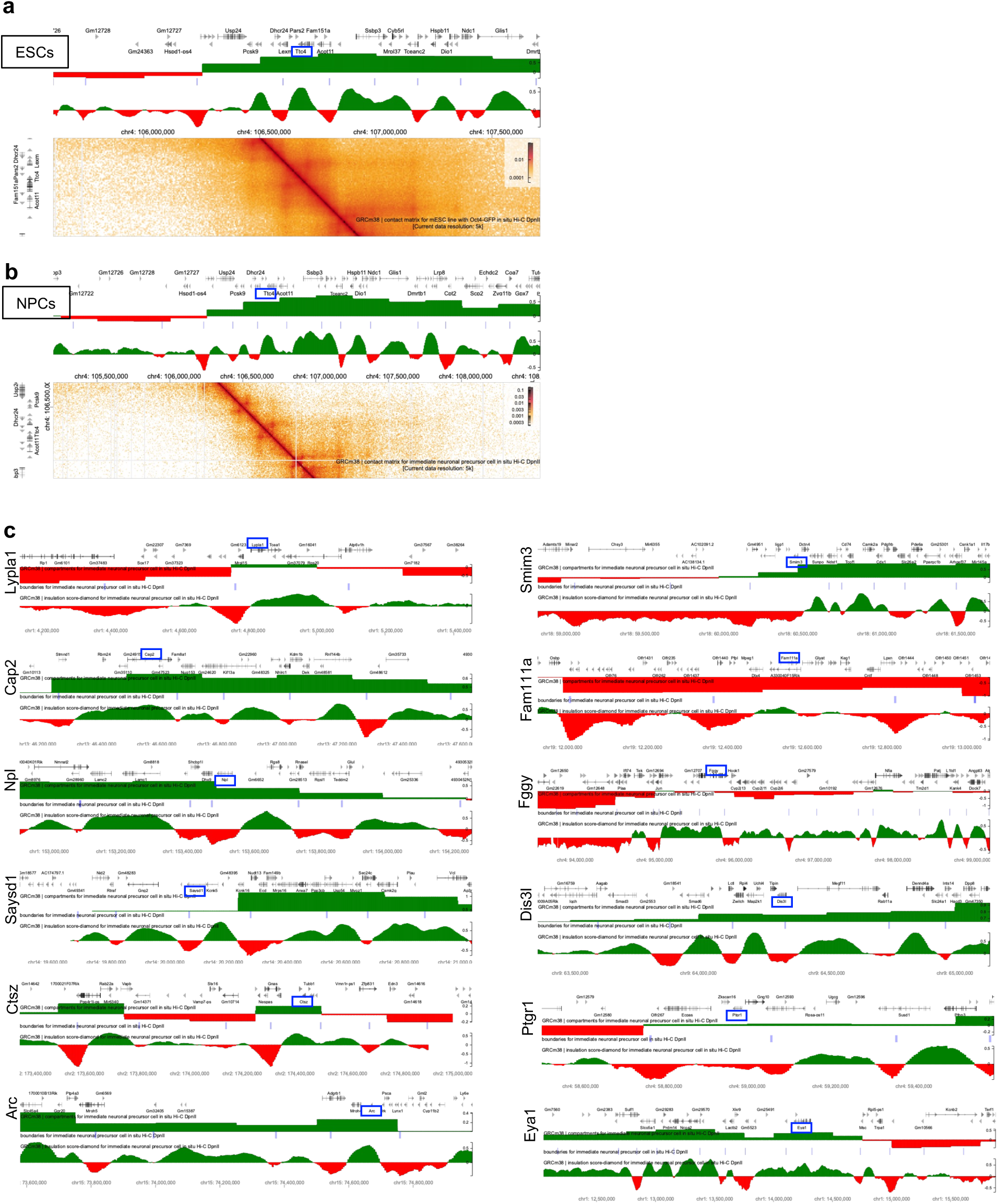
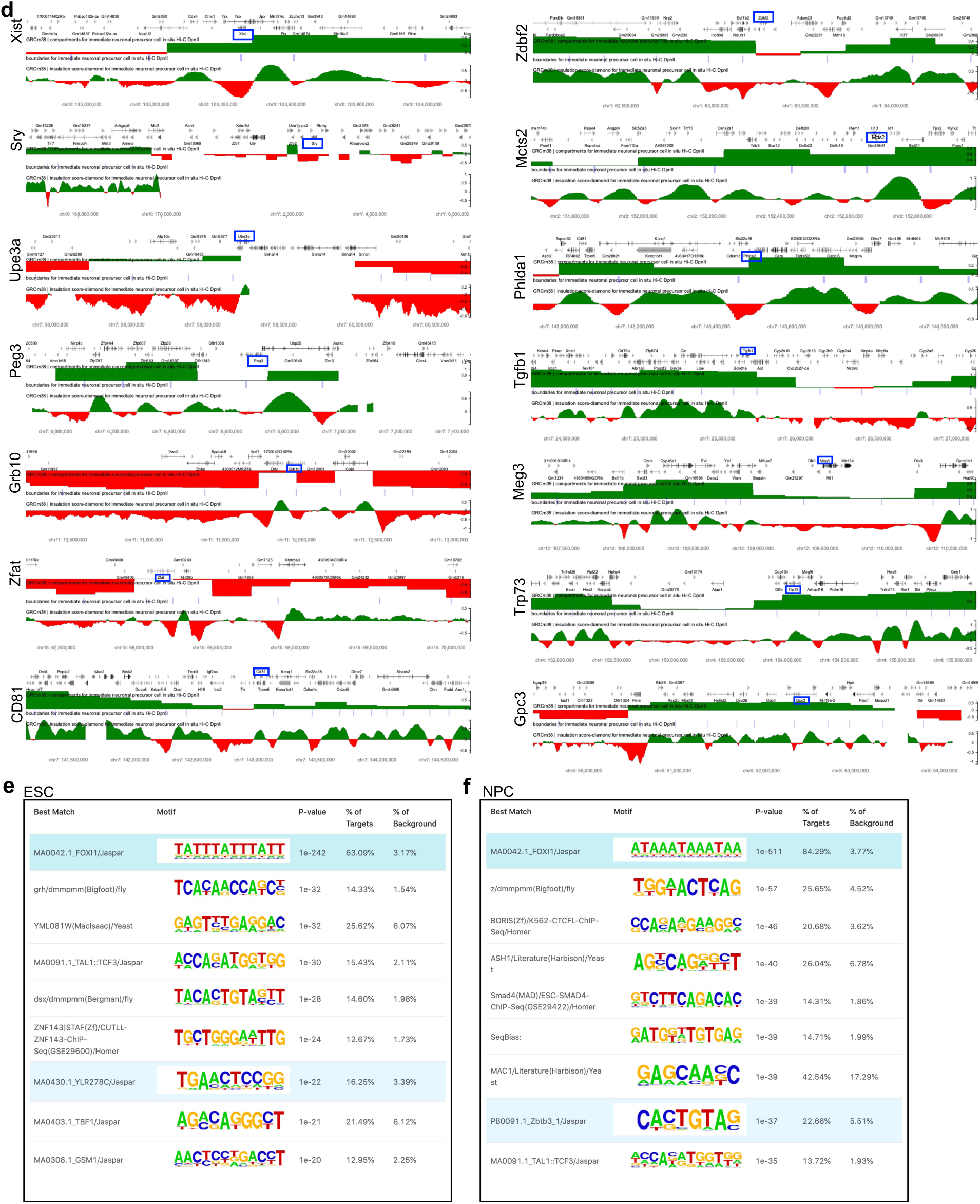
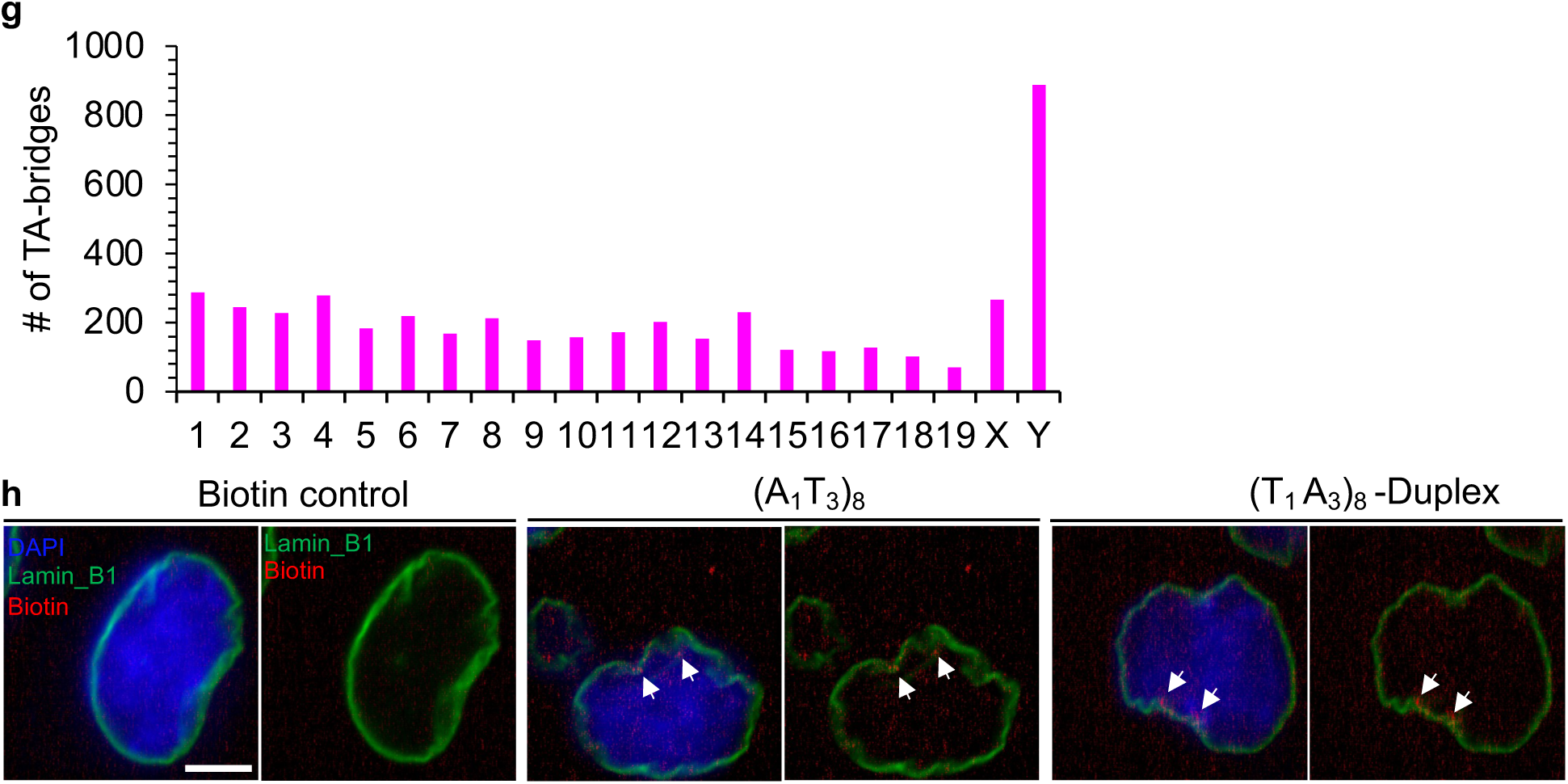
TA-bridge associated monoallelic gene’s positioning in micro-TADs within the B chromatin compartment. **a**, & **b**, *Ttc4* positioning in Hi-C-defined “micro-TADS”. The *Ttc4* gene is located in the A compartment (green). Monoallelic genes are enriched within the *Ttc4*-associated A compartment that lies within the surrounding B compartment. The contact Matrix (lower panels) for each gene shows the broader TADs and the “micro-TADs” within TADs. Vertical lines show the boundaries of the TADs. Data is from references 59, 60, 61 and 62. **c** & **d**, Association of DeMA and RaMA genes with “micro-TADs”. Panel **c** shows RaMA genes; panel **d** shows DeMA genes. **e** & **f**, De novo motif discovery. Narrow-peak search criteria were used to identify the shortest possible motifs. Only the nine most significant motifs identified in Sun1-ChIP chromatin in ESCs (**e**) and NPCs (**f**) are listed. **g**, Number of TA-bridges per chromosome. The Y chromosome contains the highest number of TA-bridges. **h**, Localization of single-stranded (A_1_T_3_)_8_ DNA and double-stranded (T_1_A_3_)_8_ DNA in NPC nuclei. At 72 hrs post-transfection, cells were fixed and immunolabeled for biotin, Lamin_B1 and DNA (DAPI). Lamin_B1 marks the nuclear periphery.

### TA-tandem repeats are enriched in monoallelic genes

Chromatin loci are positioned within specific domains in the three-dimensional nuclear space. Although it has been shown that the nuclear lamina acts as a scaffold to anchor chromatin at the NP via nuclear envelope transmembrane proteins (e.g., HP1, Lap2β, cKrox, Lamin_B receptor)^61–63^, the DNA motifs that facilitate chromatin positioning at the nuclear envelope are poorly documented. Since monoallelic genes are preferentially located in Sun1-enriched territories, we then asked whether specific motif sequences are present in the chromatin associated with Sun1 protein territories. By performing motif analysis, we discovered that *T_1_A_3_* and *A_1_T_3_* tandem repeats (which we refer to as TA-bridges) are significantly (p=1e-511 and p=1e-242, respectively) enriched in Sun1 pulled-down chromatin fractions (Fig.3c, Extended Data Fig.3e,f, Extended Data Table 4). We also found that those TA-bridges were present approximately 5 kb downstream of the 3’ end and on the fourth intron of *the Cth* gene, and ∼10 kb downstream of the 3’ end and ∼100 kb upstream of the TSS (in the first intron of *the Acot11 gene*) of the *Ttc4* gene (Fig.2h). Additionally, some genes known to be expressed monoallelically were located near these TA-bridges (*Cth*- downstream: *Zranb2*; upstream: Ankrd13c, Srsf11, Lrrc40; and Ttc4-downstream: Pars2, Dhcr24; upstream: Ssbp3, Mrpl37, Cdcp2). The monoallelic gene panels examined in the RNA-FISH assay are also linked to TA-bridges, suggesting a potential mechanism by which TA-bridges may facilitate the establishment and maintenance of monoallelic gene expression within Sun1 territories.

Through genome-wide scanning, we identified 4,572 TA-bridge motifs across the entire mouse genome. Of these, 887 (19.4%) were located on the Y chromosome (Extended Data Fig.3g). Considering the comparatively smaller size of the Y chromosome and its higher enrichment of TA-bridges compared to other chromosomes (Fig.3d), as well as its predominant localisation to the A compartment (Fig.3e), we examined the position of the Y-chromosome territory within the nucleus using published LaminB1-DamID data in ESCs from the 4DN data portal^59,60^. Although the LaminB1-DamID data (4DNFI32K5L77 – from the Bas van Steensel lab, NKI, NL) indicated that the Y chromosome is localized at the NP (Fig.3e - middle panel of enlarged area), Hi-C-defined compartments^57^ showed that the Y chromosome mainly resides in the A compartment, and its RNA expression profile^64^ aligned with the active compartment profiles (Fig.3e - lower panel of enlarged area), further supporting the presence of “micro-TADs” within LADs.

### TA-bridges localize to Sun1 territories

To further investigate whether the TA-bridge motif could direct their associated chromatin to the NP, we developed an assay to visualize TA-bridge positioning within the nucleus in live cells, using immunofluorescence microscopy, and to identify the *in vivo* TA-bridge-interacting proteins (Fig.3f). We transiently transfected biotin-labeled single-stranded (*T_1_A_3_*)*_8_* and (*A_1_T_3_*)*_8_* DNA oligonucleotides, as well as double-stranded *(T_1_A_3_)_8_* DNA into mESCs and NPCs, and observed that biotin-labeled single-stranded *(T_1_A_3_)_8_* and *(A_1_T_3_)_8_* DNA, as well as double-stranded *(T_1_A_3_)_8_* DNA, were located at the NP among the nuclear-localized signals in ESCs and NPCs (Fig.3g,h and Extended Data Fig.3h).

Next, we aimed to identify the protein candidates interacting with TA-bridges by performing mass spectrometry (MS) analysis of the protein pull-down by AT-bridges. Although previously known Sun1-binding proteins (Lmnb1, Lmnb2, Emerin) were pulled down by TA-bridges, Sun1 was not significantly enriched in the (*T_1_A_3_*)*_8_* or double-stranded TA-bridge DNA motif pull-down protein fractions (Fig. 4a, b, and Extended Data Table 5), and was also not pulled down by the (*A_1_T_3_*)*_8_* motif (Extended Data Fig. 5a). Therefore, we could not conclude that Sun1 is the direct binding protein of the TA-bridges. To further investigate the “molecular hook” that binds to the double-stranded TA-bridges, we conducted an electrophoretic mobility shift assay (EMSA) using nuclear protein extracted from mouse ESCs. In the EMSA, we observed multiple shifted biotin-positive bands for (*T_1_A_3_*)*_8_* and (*A_1_T_3_*)*_8_,* as these DNA probes most likely interacted with multiple protein complexes (Fig. 4c). Interestingly, although we noted that the double-stranded TA-bridge DNA motifs were localised at the Sun1 territory in the NP of ESCs and NPCs in the immunofluorescence assay, these motifs did not show any binding with nuclear proteins.

**Figure 4.**
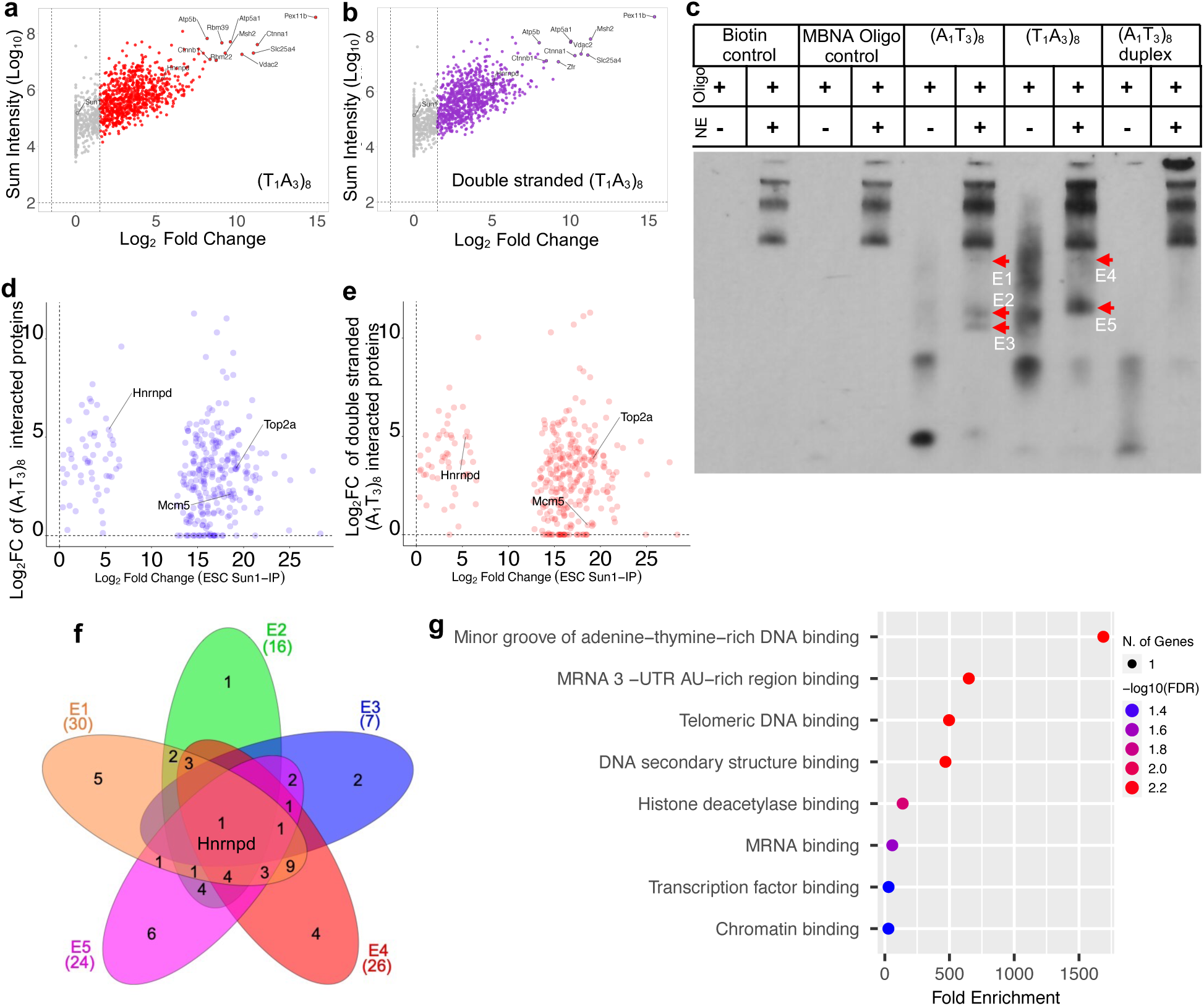
Hnrnpd interacting TA-bridges localize to Sun1 territories. **a** & **b**, Volcano plots showing in vivo interacting protein pulled down (Fig.3 f) with the single-stranded (T_1_A_3_)_8_ (**a**) and double-stranded (T_1_A_3_)_8_ (**b**) DNA motifs. Sun1 protein was detected in pull-down with both single-stranded and double-stranded (T_1_A_3_)_8_ DNA motifs; however, Sun1 was not significantly enriched. Log_10_2 of summed intensity and Log_10_1.5 of fold change were used as cutoff values for significant enrichment. **c.** EMSA assay for (T_1_A_3_)_8_, (A_1_T_3_)_8_, and double-stranded (T_1_A_3_)_8_ bridges. ESC nuclear extract was used. MBNA oligonucleotide supplied with the EMSA kit was used as the negative control. Proteins were extracted from three excised shifted bands in the (A_1_T_3_)_8_ lane and from two shifted bands in the double-stranded (T_1_A_3_)_8_ lane. Higher-molecular-weight biotin-positive bands in all samples represent non-specific interactions. **d** & **e**, Comparisons of significantly enriched proteins pulled down with Sun1, single-stranded (T_1_A_3_)_8_ DNA, and double-stranded (T_1_A_3_)_8_ DNA. Each dot represents a protein. Hnrnpd shows similar enrichment for single-stranded (T_1_A_3_)_8_ DNA. Each dot represents a protein. Hnrnpd shows similar enrichment for single-stranded (T_1_A_3_)_8_ DNA (**d**) and double-stranded (T_1_A_3_)_8_ DNA (**e**). **f**, Venn diagram showing the intersection of the most significantly enriched protein clusters obtained from five bands excised from the EMSA assay blot. Hnrnpd is the only protein common to all excised bands. **g**, Gene Ontology (molecular function) annotation for Hnrnpd. A cutoff value of FDR 0.05 was used as the significance threshold.

**Extended Data Figure 4.**
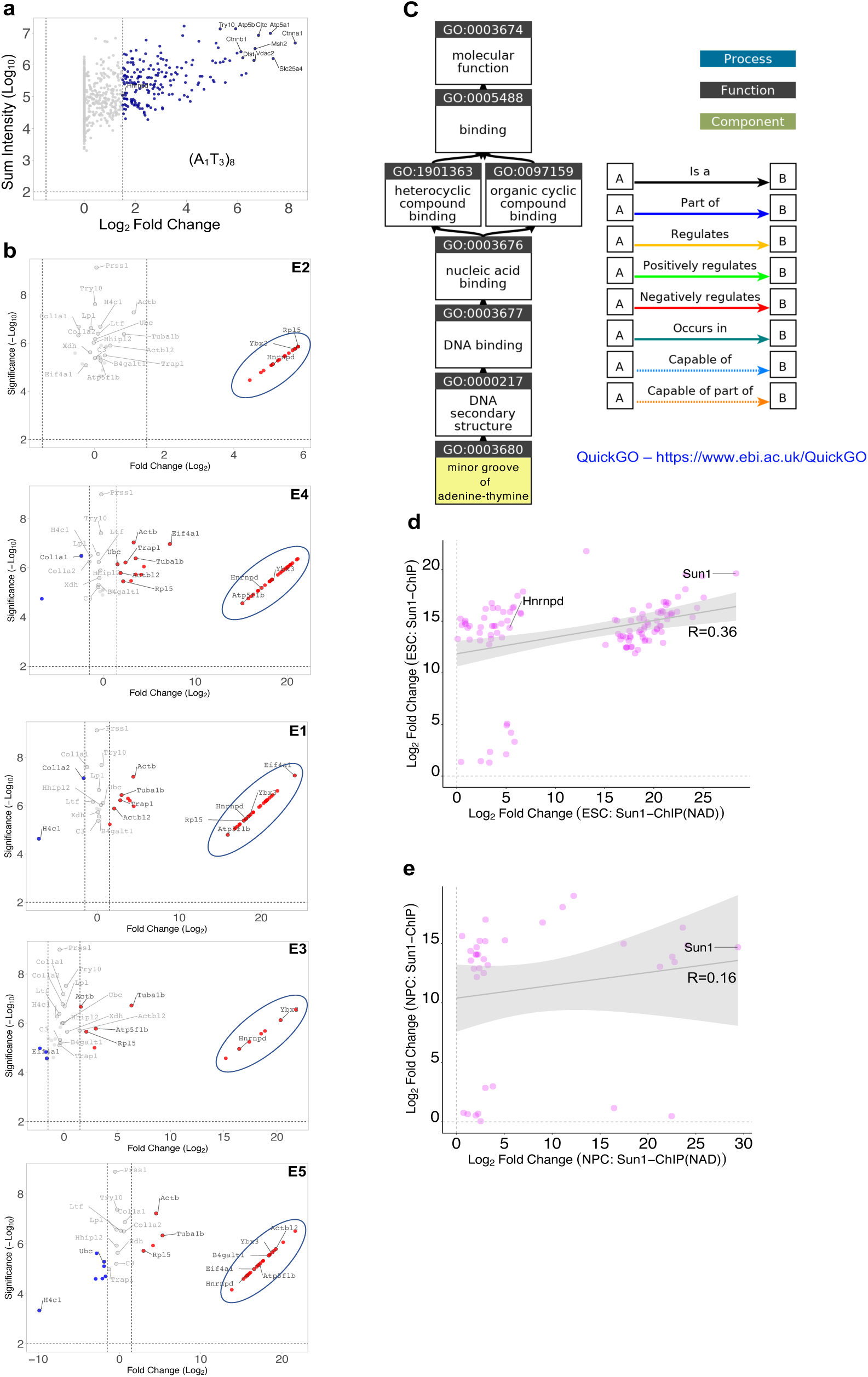
Hnrnpd interacting TA-bridges localize to Sun1 territories. **a**, Volcano plot showing significantly enriched in vivo-interacting proteins pulled down with the (A_1_T_3_)_8_ single-stranded DNA motif. Sun1 protein was not pulled down by (A_1_T_3_)_8_ single-stranded DNA motifs. Log_10_2 of summed intensity and Log_10_1.5 of fold change were used as cutoff values for significant enrichment. **b**, Volcano plots showing significantly enriched proteins identified from five biotin-positive shifted bands (E1-E5) excised from the EMSA blot. Hnrnpd is present in all highly significant protein clusters (circled). **c**, Diagram showing functional Gene Ontology annotation of Hnrnpd using QuickGO. GO term 0003680 indicates that Hnrnpd binds to the adenine-thymine-rich minor groove of DNA. **d** & **e**, Correlations analysis of significantly enriched proteins pulled down with Sun1 from non-nucleic-acid-depleted (**d**) and nucleic-acid-depleted (**e**) nuclear extracts in ESCs and NPCs. Hnrnpd is not enriched in nucleic-acid-depleted Sun1 pull-protein fractions. For ESC, the overall Pearson’s correlation coefficient (R) is 0.359 with a 95% confidence interval from -0.186 to 0.511 and a p-value of 1e-4. For NPC, the overall Pearson’s correlation coefficient (R) is 0.156 with a 95% confidence interval from -0.182 to 0.461 and a p-value of 0.364.

### Hnrnpd binds to TA-bridges

Although we identified several protein candidates known to interact with single-stranded DNA in TA-bridge MS data, to identify proteins that directly bind to TA-bridges, we performed MS analysis of proteins extracted from the multiple shifted bands identified in the EMSA assay. MS data from the five excised bands showed no enrichment of Sun1 protein (Extended Data Fig.4b and Extended Data Table 6), confirming that Sun1 does not directly interact with TA-bridges. However, by intersecting the most significant protein (Extended Data Fig.4b – circled clusters) candidates across multiple bands from the EMSA assay, we identified Hnrnpd as the only common protein candidate, which was also enriched in proteins pulled down by Sun1 (Fig.2d). To further investigate the direct interaction between Hnrnpd and TA-bridges, we examined known functions attributed to Hnrnpd. Analogous to AT-rich interacting domain (ARID)-containing proteins, Hnrnpd has been reported to bind to the adenine-thymine-rich minor grooves of B-form DNA^65^ (GO:0003680) (Extended Data Fig.4c), supporting our identification of Hnrnpd as a “molecular hook” that binds TA-bridge tandem repeats. Furthermore, our findings suggest that Hnrnpd could bind to both the noncanonical 5’-(*T_1_A_3_*)*_n_*-3’ and the 5’-*(A_1_T_3_)_n_*-3’ tandem repeat sequences. To investigate whether Hnrnpd binds to TA-bridges before or after their localisation to Sun1-enriched territories, we performed co-immunoprecipitation of Sun1-associated proteins using Sun1 antibody on nucleic acid-depleted ESC and NPC nuclear extracts, followed by MS analysis. Evaluation of the proteins immunoprecipitated with Sun1 revealed that Hnrnpd was less abundant in the Sun1 pull-down from ESC nuclear extracts (Fig.4d) and could not be detected in the Sun1 pull-down from NPC nuclear extracts (Fig.4e and Extended Data Table 7), suggesting Hnrnpd most likely binds to the AT-bridge motifs prior to the allele’s localisation to Sun1-enriched territories.

## Discussion

Although allele pairing has been observed for decades, the presence of this phenomenon in mammals has not been extensively studied^38^. Furthermore, it has not been previously demonstrated that allele pairing is necessary for some genes to be expressed from both alleles. Using the *Cth* and *Ttc4* genes, which are biallelically expressed in mESCs but become monoallelic upon differentiation, we showed that allele pairing for certain genes coincides with transcription from both alleles. In this study, we also demonstrated that for the monoallelic *Cth* and *Ttc4* genes, when the alleles are positioned further apart, only the allele associated with the Sun1-enriched domain is transcriptionally active. The establishment of *monoallelic expression, in terms of gene imprinting,* is described as a mechanism to regulate the transcriptional dosage of dosage-sensitive genes during early development, but detailed studies evaluating whether imprinted genes can be expressed biallelically in differentiated cells or in rare diseases are lacking. Although the *Cth* and *Ttc4* genes attain monoallelic status in differentiated cells, their previous biallelic expression, distinct from imprinted genes, suggests a degree of plasticity in the transcriptional status of the inactive allele. Consequently, reactivation of the silent allele—particularly in the context of “disease causative monoallelic genes”—may have clinical significance. Additionally, our finding that the active alleles of monoallelic genes are localised to Sun1-enriched territories at the NP suggests that their gene expression may be facilitated within these specialised nuclear compartments.

Our observation of monoallelic genes’ preferential expression at the NP supports the idea that the chromatin located at the NP is not solely heterochromatic^66–70^. Evaluating published data, we also demonstrated that the active alleles of monoallelic genes are cluster-embedded within the B compartment at the NP. However, why many monoallelic genes adopt this unique arrangement at the NP remains enigmatic. In our previous studies, we showed that most monoallelic genes encode only one RNA isoform, which is translated into a functional protein^22^. . Therefore, it is reasonable to argue that NP positioning may assist RNAs in being exported through nuclear pore complexes to avoid the high rate of RNA turnover in the nucleus— acting as a “defensive mechanism,” as proposed in the gene gating hypothesis, an idea worth exploring further. Moreover, since the transcriptionally active alleles are localized at the NP, these genes (monoallelic: Rac1, S100a10, Junb; biallelic: Sun1, Scn5a) may be at risk of misregulation in diseases caused by laminopathy and nuclear deformation^71^, and thus could have clinical relevance.

Although in association with SPDYA, SHELTERIN, and TTM complex proteins, Sun1 stabilizes the telomeric rings at the inner nuclear membrane^72^, Sun1 does not serve as a direct binding partner to the DNA. Therefore, in line with our findings, it is likely part of a scaffold that harbors chromatin. The discovery of novel repetitive elements in genomes has been achieved using long-read DNA sequencing methods^73,74^. However, so far, only a few repetitive elements have been linked to genome regulation^75–78^, and the functional significance of many remains elusive. Although some transcriptionally active genes have been reported to be localized at the NP^66–70^, the mechanism by which alleles are physically positioned and maintained at the NP remains unknown. Here, we discovered that Hnrnpd physically interacts with TA-bridges, highlighting the potential importance of other yet-to-be-identified DNA tandem repeat elements. While Hnrnpd binds to TA-bridges, it is not an enriched component of Sun1 territories, suggesting that Hnrnpd’s binding to the TA-bridges occurs before the localisation of alleles to the Sun1 territories. Furthermore, we observed that known helicases (Mcm5^79^ and Top2a^80^) are significantly enriched among TA-bridge interacting proteins (Fig.4d,e), indicating that strand separation of the TA-bridge likely initiates prior to the motifs being localised at the Sun1 nuclear territories.

During stem cell differentiation into more specialized cells, the gene expression program changes alongside chromatin reorganisation, leading to different architectures depending on the progenies^81,82^. Furthermore, it has been shown that asynchronous allele replication occurs more frequently during cell differentiation, potentially promoting the early-replicating allele to become the active allele of a monoallelic gene^83,84^. Because Hnrnpd binds to TA-bridges independently of Sun1, we propose a mechanistic model where Hnrnpd attaches to single-stranded TA-bridges of the early replicating allele in prophase, resulting in the targeting of the TA-bridge-bound Hnrnpd to the newly forming nuclear membrane Sun1-enriched domains in telophase via a molecular targeting mechanism to be elucidated in future studies.

## Methods

### Mouse ESC (mESC) culture and *in vitro* neural progenitor cell (NPC) differentiation

Polymorphic F1 mESCs were generated and maintained as previously described^22^. NPC in vitro differentiation was carried out as follows. Male F1 mESCs were seeded on 0.1% gelatin-coated 6-well plates at a density of 1.0 x10^5^ cells per well in 2.5 ml of 2i/LIF medium. At 24 hrs post-seeding, cells were washed three times with 1x PBS and switched to NDiff^®^ 227 neural induction medium (TaKaRa, Cat.No. Y40002). Cells were incubated at 37^0^C, 5% CO_2_. Approximately 48 hrs after neural induction, mESCs changed their morphology to neural-like cells. At this point, cells were washed three times with 1x PBS, and 2.5 ml of NPC maintenance medium supplemented with 20 ng/ml murine EGF (PeproTech Inc, Cat.No. 315-09) and 20 ng/ml murine Fgf2 (PeproTech Inc, Cat.No.450-33) was added to each well and cultures were incubated for an additional 36 hrs. Subsequently, cells were trypsinised and reseeded at a density of 1x10^5^ cells per well in NPC maintenance medium to establish long-term NPC cultures.

### Establishment of an allele-specific gene expression visualization system

Generating a parental mESC line with mCherry-PP7CP and EYFP-MS2CP Tet-On inducible expression system: The Tet-On Inducible Expression System® plasmid (Clontech, Cat. No. 634301) was linearized by double digestion with EcoRI and AgeI. The mCherry-PP7CP DNA fragment was excised from the PP7-mCherry plasmid (Spector lab) by double digestion with BaeI, and adaptors compatible with the linearized Tet-On vector were ligated to the 5’ end (annealed Oligo#1 + Oligo#2) and to the 3’ end (annealed Oligo#3 + Oligo#4), generating the Tet-On-PP7CP-mCherry plasmid. The EYFP-MS2CP fragment was excised from the pMS2-cp-YFP plasmid (Spector lab) using NheI and NgoMIV restriction enzymes, and adaptors compatible with the linearized Tet-On vector (annealed Oligo#5 + Oligo#6 at the 3’ end) were ligated to generate the Tet-On-MS2CP-EYFP plasmid. Both Tet-On-PP7CP-mCherry and Tet-On-MS2CP-EYFP plasmids were double-digested with BglII and PclI to excise the expression cassettes. Subsequently, annealed Adpt A (Oligo#7 + Oligo#8) and annealed Adpt B (Oligo#9 + Oligo#10) were ligated into the mCherry end of the excised expression cassette and the Tet-On-3G end of the Tet-On-PP7CP-mCherry plasmid, respectively. Likewise, Adpt C (Oligo#11 + Oligo#12) and annealed Adpt D (Oligo#13 + Oligo#14) were ligated into the EYFP end of the excised MS2CP expression cassette and the Tet-On-3G end of the Tet-On-MS2CP-EYFP plasmid, respectively. The resulting expression cassettes, constructed with overhanging sequences complementary to the insertion site at the Rosa26 locus, were used for transfection into mESCs.

Oligos:

Oligo#1-CCGAGTGAGTCGTATTA

Oligo#2-TCGATAATACGACTCACT

Oligo#3-AAGCCAAAAAAGAAGAGAAAGGTATAA

Oligo#4-AATTTTATACCTTTCTCTTCTTTTTTGGCTTGGACA

Oligo#5-CTAGTAATACGACTCACTATAGG

Oligo#6-AATTCCTATAGTGAGTCGTATTA

Oligo#7-GATCGGATGGATCCATCGGAAGGGGAGCCTTTCTCTCTGGGCAAGAGCGGTGCAATGGTGTGTAAAGGTAGCCGCTC

Oligo#8-CGATGGATCCATCC

Oligo#9- GGATGGATCCACTG

Oligo#10-CATGCAGTGGATCCATCCTCTCGGCTACCTTTACACACCATTGCACCGCTCTTGCCCAGAGAGAAAGGCTCCCCTTC

Oligo#11-GATCGGATGGATCCATCGTCTGAAGGAGAGCCTTTC

Oligo#12- CGATGGATCCATCC

Oligo#13-CGATGGATCCATCC

Oligo #14-CATGCAGTGGATCCATCCGAAAGGCTC**T**CCTTCATC

Designing allele-specific gRNAs for expression cassette knock-in at the Rosa26 locus: Allele-specific gRNAs were designed to incorporate SNPs in the Rosa26 (R26) alleles at the PAM site and were cloned into pSpCas9n(BB) (PX460) (Addgene, Cat.No. 48873) plasmid. The following gRNA sequences were used in combination. gR26#1 (annealed R26-5’Cast1.1 and R26-5’Cast1.2), gR26#2 (annealed R26-3’Cast-Comp1.1 and R26-3’Cast-Comp 1.2), gR26#3 (annealed R26-5’B6 1.1 and R26-5’B61.2), gR26#4 (annealed R26-3’B61.1 and R26-3’B61.2). Next, the indicated combinations of gRNAs and adaptor-ligated expression cassettes were transfected into F1 mESCs using the Xfect transfection kit (Takara, Cat.No.631317). To target the Tet-On-PP7CP-mCherry expression cassette to the R26 CAST (paternal) allele, gR26#1, gR26#2 and Tet-On-PP7CP-mCherry cassettes were co-transfected. To targt the Tet-On-MS2CP-EYFP to the R26 C57 (maternal) allele, gRNA#3, gRNA#4, and the Tet-On-MS2CP-EYFP were co-transfected. For each transfection, 2 μg of gRNA-Cas9 plasmids and 1.0 μg of adaptor-ligated cassette DNA were used. Prior to transfection, mESCs were seeded at 0.5x10^5^ cell/ well in 6-well Corning tissue culture plates (Corning, Cat. No. 353046) and cultured for 12 hrs. At 24 hours post-transfection, cells were rinsed with KnockOut™ DMEM (Gibco, Cat. No. 10829-018) containing 10% FBS, then switched to fresh 2i medium. At 48 hours post-transfection, single clones were picked under a microscope and expanded.

Oligos:

R26-5’Cast1.1-CACCGAGTGGGGTCGACTATCTGAAGGG

R26-5’Cast1.2-AAACCCCTTCAGATAGTCGACCCCACTC

R26-3’Cast-Comp1.1-CACCGGCTTGCCCTTTTCGTCTTCTCGG

R26-3’Cast-Comp1.2-AAACCCGAGAAGACGAAAAGGGCAAGCC

R26-5’B61.1-CACCGTAGTGGGGTCGACTATCTGAAGG

R26-5’B61.2-AAACCCTTCAGATAGTCGACCCCACTAC

R26-3’B61.1-CACCGCCGCTCTTGCCCAGAGAGAAAGG

R26-3’B61.2-AAACCCTTTCTCTCTGGGCAAGAGCGGC

MS2 and PP7 stem-loops construction: The pCR4-24xMS2SL (Addgene, Cat. No.31865) and pCR-24xPP7SL (Addgene, Cat. No. 31864) plasmids were double-digested using SpeI and NotI, and the 24xMS2SL (1352bp) and 24xPP7SL (1492bp) DNA fragments were gel-purified. Annealed adapters were ligated into each of the excised stem-loop DNA fragments as follows: Ttc4 gene C57a allele (Lig#1): annealed Ttc4-B6-5.1 + Ttc4-B6-5.2 and annealed Ttc4-B6-3.1 + Ttc4-B6-3.2.

Ttc4 gene CAST allele (Lig#2): annealed Ttc4-CAST-5.1 + Ttc4-CAST -5.2 and annealed Ttc4-CAST -3.1 + Ttc4-CAST -3.2.

Cth gene C57 allele (Lig#3): annealed Cth-B6-5.1 + Cth-B6-5.2 and annealed Cth-B6-3.1 + Cth-B6-3.2.

Cth gene CAST allele (Lig#5): annealed Cth-CAST-5.3 + Cth-CAST-5.4 and annealed Cth-CAST-3.3 + Cth-CAST-3.4.

Oligos:

Ttc4-B6-5.1 TAATACGACTCACTATAGG

Ttc4-B6-5.2

GGCCCCTATAGTGAGTCGTATTAAAAGGGAAAACAACACAAGCCTACTTGCAAACCACTGCCCATTGACTCCTCAAGCTTCTGTCAGATGTGGCACAAGCCCTTGCATGGGCATTGAGAAGTAGAGAATCCAGTAGGGCTCTGAGTAGTCCCAGGATCCAAGT

Ttc4-B6-3.1 CTAGTAATACGACTCACTATAGGACTTGGATCCTGGGACTACTCAGAGCCCTACTGGATTCTCTACTTCTCAATGCCCATGCAAGGGCTTGTGCCACATCTGACAGAAGCTTGAGGAGTCAATGGGCAGTGGTTTGCAAGTAGGCTTGTGTTGTTTTCCCTTT

Ttc4-B6-3.2 CCTATAGTGAGTCGTATTA

Ttc4-CAST-5.1 AATTAACCCTCACTAAAGGG

Ttc4-CAST-5.2 CTAGCCCTTTAGTGAGGGTTAATTACAGGATGTGGCTTCATGGCTCC AGTC

Ttc4-CAST-3.1 GGCCAATTAACCCTCACTAAAGGGACTTGGATCCTGGGACTACTCAGAGCCCTACTGGATTCTCTACTTCTCAATGCCCATGCAAGGGCTTGTGCCACATCTGACAGAAGCTTGAGGAGTCAATGGGCAGTGGTTTGCAAGTAGGCTTGTGTTGTTTTCCCTTTATCGTGAGGCCTCAT

Ttc4-CAST-3.2 CCCTTTAGTGAGGGTTAATT

Cth-B6-5.1 AGAACTGTCTTCTGTTTATCTTCTAACTAACAGGTTGTTCTGTTAGTATCATTTCGGTAATTTTGCTATATTTGTGTCCAAGGAAGTAAGAGTTGTTCTGTAATACGACTCACTATAGG

Cth-B6-5.2 GGCCCCTATAGTGAGTCGTATTA Cth-B6-3.1 CTATAGTGAGTCGTATTAGATC

Cth-B6-3.2 TAATACGACTCACTATAGGTTCATTCTCAACAAGACAAAACCCTAGTACAACAGAAAAAAGAAAGAAGTCATCGAATACTATATAAAATTAGTACAAATGTTACAATGTTAAATCATAACTACAAAATACTTCAATTTAATAATTTACTTACCAGAATTTAG GTGAC

Cth-CAST-5.3 AATTAACCCTCACTAAAGGG

Cth-CAST-5.4 CTAGCCCTTTAGTGAGGGTTAATTCCTTGGACACAAATATAGCAAAATTACCGAAATGATACTAACAGAACAACCTGTTAGTTAGAAGATAAACAGAAGACAGTCCTTAA

Cth-CAST-3.3 GGCCAATTAACCCTCACTAAAGGGTAAAGGTTGTACTTGATACTTGTTATTTTTCTTAAATAAATCCAGTT

Cth-CAST-3.4 CCCTTTAGTGAGGGTTAATT

gRNA cloning for allele-specific insertion of stem-loops: To clone gRNAs into pSpCas9n(BB) (PX460) plasmids, we generate plasmid libraries by cloning allele-specific gRNA pairs as described below.

T8: Ttc4-C57BL-gRNA-1.1: + Ttc4-C57BL-gRNA-1.2:

T9: Ttc4-C57BL-gRNA-2.1: + Ttc4-C57BL-gRNA-2.2:

T10: Ttc4-CAST-gRNA-3.1: + Ttc4-CAST-gRNA-3.2:

**T11**: Ttc4-CAST-gRNA-3.1a: + Ttc4-CAST-gRNA-3.2a:

T12: Ttc4-CAST-gRNA-4.1a: + Ttc4-CAST-gRNA-4.2a:

T13: Ttc4-CAST-gRNA-4.1: + Ttc4-CAST-gRNA-4.2:

C1: Cth-C57BL-gRNA-1.1: + Cth-C57BL-gRNA-1.2:

C2: Cth-C57BL-gRNA-2.1: + Cth-C57BL-gRNA-2.2:

C3: Cth-CAST-gRNA-3.1: + Cth-CAST-gRNA-3.2:

C4: Cth-CAST-gRNA-3.1a: + Cth-CAST-gRNA-3.2a:

C5: Cth-CAST-gRNA-4.1a: + Cth-CAST-gRNA-4.2a:

C6: Cth-CAST-gRNA-4.1b: + Cth-CAST-gRNA-4.2b:

C7: Cth-CAST-gRNA-4.1: + Cth-CAST-gRNA-4.2:

Ttc4-C57BL-gRNA-1.1: CACCGTTCTGTTGACTGTATAAACTTGG

Ttc4-C57BL-gRNA-1.2: CAAGACAACTGACATATTTGAACCCAAA

Ttc4-C57BL-gRNA-2.1: CACCGAAATGAGACCTCACAATAAAGGG

Ttc4-C57BL-gRNA-2.2: CTTTACTCTGGAGTGTTATTTCCCCAAA

Ttc4-CAST-gRNA-3.1: CACCGTGTGGATGCCATGAAGTGACTGG

Ttc4-CAST-gRNA-3.2: CACACCTACGGTACTTCACTGACCCAAA

Ttc4-CAST-gRNA-3.1a: CACACCTACGGTACTTCACTGACCCAAA

Ttc4-CAST-gRNA-3.2a: CAAGACAACTGATATATTTGAACCCAAA

Ttc4-CAST-gRNA-4.1a: CACCGGTTTATATAGTCAACAGAACAGG

Ttc4-CAST-gRNA-4.2a: CCAAATATATCAGTTGTCTTGTCCCAAA

Ttc4-CAST-gRNA-4.1: CACCGCTACTCCTATTTCAGAAATGAGG

Ttc4-CAST-gRNA-4.2: CGATGAGGATAAAGTCTTTACTCCCAAA

Cth-C57BL-gRNA-1.1: CACCGAAACAGAAGACAGTTCTTAAGGG

Cth-C57BL-gRNA-1.2: AAACCCCTTAAGAACTGTCTTCTGTTTC

Cth-C57BL-gRNA-2.1: CACCGTGGTCTTAAATCCACTGTGGTGG

Cth-C57BL-gRNA-2.2: AAACCCACCACAGTGGATTTAAGACCAC

Cth-CAST-gRNA-3.1: CACCGCTTCCATGGATCTTCCCTTAAGG

Cth-CAST-gRNA-3.2: AAACCCTTAAGGGAAGATCCATGGAAGC

Cth-CAST-gRNA-3.1a: 806 CACCGCAGAACAACTCTTACTTCCTTGG

Cth-CAST-gRNA-3.2a: AAACCCAAGGAAGTAAGAGTTGTTCTGC

Cth-CAST-gRNA-4.1a: CACCGAGTAAGAGTTGTTCTGTTTTGGG

Cth-CAST-gRNA-4.2a: AAACCCCAAAACAGAACAACTCTTACTC

Cth-CAST-gRNA-4.1b: CACCGAAACCACCACAGTGGATTTAAGG

Cth-CAST-gRNA-4.2b: AAACCCTTAAATCCACTGTGGTGGTTTC

Cth-CAST-gRNA-4.1: CACCGAAGAAAAATAACTTGATAACTGG

Cth-CAST-gRNA-4.2: AAACCCAGTTATCAAGTTATTTTTCTTC

Allele-specific MS2 and PP7 stem-loop knock-in to the *Cth* and *Ttc4* genes: A single-cell-derived mESC clone expressing mCherry-PP7CP and EYFP-MS2CP was transfected with a mixture of DNA constructs and gRNA-Cas9 plasmids using the Xfect transfection reagent to knock in stem loops to individual alleles. Prior to transfection, 0.5x10^5^ mESCs per well were plated in 6-well plates and cultured for 12 hours. To target the *Cth* gene alleles, Lig#3 and Lig#5 (3.45 μg of DNA in each) were co-transfected with gRNAs C1, C2, C3, C4, C5, C6 and C7 (0.3 μg of plasmid each). To target the *Ttc4* gene alleles, Lig#1 and Lig#2 (3.6 μg of DNA each) were co-transfected with gRNAs T8, T9, T10, T11, T12 and T13 (0.3 μg of plasmid each). Twelve hours after transfection, the medium was replaced with fresh 2i medium. At 24 hrs post-transfection, single colonies were picked under a light microscope, expanded and genotyped using Cth-Wt gene F (AAAGTTCGAGTVAAAGCTGTCAT and Cth-Wt gene R (TCAAGTACAACCTTTAATCTCTTGAA) for Cth clones, and Ttc4-Wt gene F (GGCAGTGGTTTGCAAGTAGG) and Ttc4-Wt gene R (CCAGCTGGTTTCAGTTTAATAATGT) for *Ttc4* clones. The mESC clones for both *Cth* and *Ttc4* were further validated for on-target integration by DNA-FISH assay.

### SDS-polyacrylamide gel electrophoresis and immunoblotting

Protein electrophoresis was performed using 1.0 mm-thick and 8x10 cm Tris-glycine SDS-polyacrylamide gels cast with 5% Stacking gel (30% acrylamide mix, 1.0 M Tris (pH6.8), 10% SDS, 10% ammonium persulfate, 0.1% TEMED in dH_2_O) and 12 % resolving gel (30% acrylamide mix, 1.5 M Tris (pH8.8), 10% SDS, 10% ammonium persulfate, 0.1% TEMED in dH_2_O). Gels were run in 1x Tris-glycine SDS running buffer, and proteins were transferred in ice-cold 1x transfer buffer onto Immobilon^®^-P Transfer membranes (Merck Millipore, Cat.No. IPVH00005). The transfer membranes were air-dried to dehydrate and stored until use. For immunoblotting, membranes were rehydrated in 0.1% PBST (0.1% Tween-20 in 1xPBS), changing the buffer several times until the membranes were fully wetted. Rehydrated membranes were blocked for 1 ½ hours in 5% blocking buffer containing skim milk (Sigma, Cat.No. 1153630500) prepared in 0.1% PBST and then incubated overnight with primary antibodies diluted in the same blocking buffer. The next day, membranes were washed four times in 0.1% PBST for 10 minutes each and incubated for 2 hours at room temperature with HRP-conjugated secondary antibody buffer (3% milk in 0.1% PBSTX). After the incubation, membranes were washed four times for 10 minutes each with 0.1% PBST, followed by a final wash in 1x PBS. ECL reagent (Thermo Fisher, Cat. No. PI32209) was added to fully cover the membrane, and chemiluminescent signals were captured on X-ray film (Amersham Hyperfilm™ ECL, GE Healthcare Limited, Cat.No. 28906836). Primary antibodies used: Goat anti-YFP polyclonal antibody (1:500) (ORIGENE, Cat.No. AB1166-100); mouse anti-mCherry antibody (1:500) (Thermo Fisher, Cat.No. MA5-32977); Histone H3 antibody (1:1000) (Novus Biological, Cat. No. NB500-171-0.025 ml). Secondary antibodies: Donkey anti-Goat IgG (H+L), HRP-conjugated (1:500) (Thermo Fisher, Cat.No. A15999); Donkey anti-Mouse IgG (H+L), HRP-conjugated (1:500) (Abcam, Cat. No. AB205719).

### Immunofluorescence assay

Cells were cultured on ibidi u-Slide 8-well glass-bottom chambers (ibidi GmbH, Cat. No. 80827). Cells were washed once with 1x PBS and then fixed in 4% PFA for 12 minuites with gently rocking. PFA was removed by rinsing three times with 400 μl of 1x PBS. Cells were permeabilized with 0.3% PBSTx (0.3% Triton X-100 in 1xPBS) for 10 minuites at RT with gental rocking, followed by three washes with 400 μl of 0.1% PBSTx. Cells were blocked for one houre at RT in blocking buffer (3% BSA, 10% NDS, 0.1% PBSTx). Then 200 μl of blocking buffer containing primary antibodies was added to each well, and incubated overnight at 4°C on a shaker. The next day, primary antibodies were removed by washing four times with 400 μl of 0.1% PBSTx for 10 minuits each. Secondry antibodies diluted in buffer (5% NDS in 0.1% PBSTx) were added at 200 μl per well and incubated for 1 ½ hours at RT on a slow rocker. After incubation, secondary antibodies were removed by washing with 0.1% PBSTx three times, fllowed by a final wash with 1X PBS. DAPI diluted 1:10000 in 1x PBS (200 ul) was added to each well and incubated for 5 minuites, followed by three washes with 1x PBS. A total of 100 μl of ibidi mounting medium (ibidi GmbH, Cat.No. 50001) was added to each well, and chamber slides were wrapped in aluminium foil and stored at 4°C until imaging. Primary antibodies: Sun1 (1:700, gift from Min lab), anti-Biotin (1: 1000) (Santa Cruz, Cat. No. 101339), Lamin_B1 (1:500) (Abcam, Cat. No. ab16048). Secondary antibodies: Donkey anti-Rabbit AF568 (Thermo Fisher, Cat.No. A10042), Donkey anti-Mouse, AF647 (Thermo Fisher, Cat.No. A-21235), Goat anti-Rabbit AF488 (Thermo Fisher, Cat. No. A-11008). Secondary antibodies were used at 1:1000 dilution.

### Live imaging using lattice light-sheet microscopy and quantitation

mESCs were cultured overnighrt on small round NO. 1 cover glass, 5 mm diameter (Thomas Scientific, Cat.No.1217H19). Four hourse before image acquisition, medium was replaced with medium containing 20 ng/ml doxycycline (Sigma, Cat.No. 324385-1GM), and the same medium condition was used during image acquisition. Live images was performed using a 3i lattice light-sheet microscope (LLSM) with the following parameters: 62.5x LLS objective, magnification changer 1.0x, binning 1x1, average time-lapse interval 20 seconds, pixel size (XY): 0.104 μm, Z-step 0.553 μm. Channel 1: 488nm excitation at 300 ms; Channel 2: 560nm excitation at 300 ms. The system uses a 25x/ 1.1 NA detection objective, with additional optical manginification to achive a final magnification of 62.5x. Each laser was set to 9% AOTF output, with a laser head power of 50 mW each. Images were deskewed and deconvolved using the algorithms embedded in SlideBook6 software.

### Nick translated probe preparation

One microgram of PCR-generated DNA probe was used in each reaction. The reaction mixture (total 45 μl) contained DNA fragments, 3.8 μl labeled dUTP (0.2 mM), 2.5 μl of 0.1 mM dTTP, 10 μl (0.1 mM) dNTP mix, 5.0 μl of 10x Nick Translation buffer, and nuclease-free ddH_2_O (volume up to 45 μl). The mixture was vortexed and briefly spun down, followed by addition of 5.0 μl of Nick Translation enzyme. Sampels were gently vortexed and centrifuged, then incubated at 15°C for 10 hours, heat-inactivated at 70°C for 10 minutes, and cooled to 4°C. Samples were transferred to 1.5 ml RNase-free microcentrifuge tubes, and DNA was precipitated by adding 1 μl of 0.5 M EDTA, 1 μl of linear acrylamide (Ambion), 5 ul of 3M NaOAc (pH5.2), and 125 μl of cold (−20°C ) 100% ethanol, followed by mixing and incubation at -80°C for 2 hours. Samples were centrifuged at 20000 rcf for 60 minuites at 4°C to pallet DNA. The pellet was washed twice with 1 ml of 75% ethanol, centrifuged at 14000 rpm for 5 minuites each time. Pellet were dried at 37°C for 15 minuites, resuspended in 50 μl of DEPC-treated DNase/RNase free H_2_O, and incubated at 37°C on a heat block with shaking for 1 hour (at 200 rpm). Probes were stored at - 20°C until use.

### Nascent RNA-FISH assay

NPCs were cultured in 24-well plates (MatTek, Cat.No. P24G013F) and used at ∼80% confluency. All buffers were prepared using DEPC-treated 1x PBS. Freshly prepared 4% PFA (in DEPC-treated PBS) was chilled on ice, then warmed at 37°C for 30 minutes before use. Cells were rinsed once with 1x PBS, fixed with 200 μl of 4% PFA for 15 minutes at RT with gentle shaking, and washed three times with 1x PBS (10 minutes each). A 200 mM VRC (stock) was warmed at 60°C before used. A 0.5% Triton X-100/ 2 mM VRC permiabalization buffer was in 1x PBS, and 300 μl was added per well wth incubation on ice for 10 minutes. Cells were washed three times with 1x PBS, then saturated with 400 μl of 2x SSC for 2 hours at 37°C in a humidified chamber. Probe preparation: A mixture of 100 ng DNA probe and 2 μl competitor yeast tRNA (10 μg/ul) was dried in SpeedVac at maximum heat for 25 minutes. Dried probes were dissolved in 10 ul deionized formamide (Ambion), incubated at 37°C for 20 minutes, denatured at 95°C for 10 minutes, and chilled on ice for 10 min. Hybridization buffer was prepared by mixing 4 μl warmed 50% dextran sulfate (42°C), 3 μl DEPC-treated ddH_2_O, 1 μl RNasin (Promega), and 2 μl 20x SSC. Ten microliters of hybridization buffer was added to a each probe tube and mixed gently. A total of 40 μl of probe mixture was added to each well, sealed with Parafilm, wrapped in foil, and incubated for 16 hours at 37°C in a humidified incubator. After hybridization, cells were washed three times (5 minutes each) with pre-warmed (42°C) 2x SSC/ 50% deionized formamide, then sequentially washed with: pre-warmed (42°C) 2x SSC three times (5 minutes each), pre-warmed (37°C) 1x SSC three times (5 minuits each), 0.5% SSC three times (5 minutes each). Cells were then incubated with 200 μl of DAPI in 4x SSC for 5 minutes and washed once with 4x SSC. Finally, 100 μl ibidi mounting medium (ibidi GmbH, Cat.No.50001) was added per well, and plates were wrapped in aluminum foil and stored at 4°C until imaging.

Primers used for PCR amplification to generate nick translated probes:

*Acyp2*: F- GGGCCGACTTTTCTAACGA, R- TGAATGACTCCATAAAACAACATT

*Arc*: F- GTGAAGACAAGCCAGCATGA, R- CTCCAGGGTCTCCCTAGTCC

*Fggy*: F-CCTGGCATTGCAAAATATGA, R- CCTGGATCCAATGAGTGAGC

*Bag3*: F-CGAGAGCCTCCACCTGTTAC, R- CCAGACAACCAGAGGGGTTA

*Eya1*: F- TGTCAGTGCTGGACTTCAGG, R- CTGCTGCCCCCAGTATTAAA

*Dis3l*: F- CGCTGCCACTCTGATACAAT, R- TCTGAAGACAGAGGCAAGAGG

*Lypla1*: F- AGTCACCTCCTCCTGCTGAA, R- CCAGACAACCAGAGGGGTTA

*Cap2*: F- GCCTTCCCTGAGTCTCCTCT, R- GCGCTATGCTTCGTTACACA

*Cth*: F- CCTCTGTGCCTGAGAAGGAC, R- TGAATTTCACAGTAGGTATTCCAAA

*Npl*: F- CATCATGACGCTGGTCTCTG, R- CCCAAAAGGCAGGTGAAAT

*Saysd1*: F- GTCCTGCTGGGACTGTTTGT, R- TCCTCAGGCACATGGTACAC

*Smim3*: F- CTATCACCCAATCCCCTGTG, R- TGGGTTGTAGCTCAGTGGTG

*Fam111a*:F- GGGACCAAAGACCACTTTCA, R- GATCCAGAAGACCCACCAAA

*Ctsz*: F- TCGTGACCAGCACCTACAAG, R- GGATTCCAGGGCTCCAGAT

*Ptgr1*: F- TCCCTGGAGGGTTAGCTTTT, R- AATGGGCTTAGGGAATGGAC

*Cd81*: F- ATTCTGAGCATGGTGCTGTG, R- CTTGAAATGGAGGCAGGAAA

*Meg3*: F- TCAAAGGGCTGAGGAGAAAA, R- AAGTGGTCAGATTGCGAAGC

*Xist*: F- TGTGTATTGTGGGTGTGTCTATTT, R- AGTCACGAAAGGCAGGAGTG

*Gpc3*: F- CAAGCAGCATGGAAATCAGA, R- TGTCAAAGAAATCCATGCAGA

*Tssc4*: F- ATGACACCCTGCCTTCTGAC, R- CACACCTTCCCACTTGGACT

*Ube3a*: F- CCACCTAGTCCTCTCGTCCA, R- CGCCTCACTGGTCCTTGT

*Sry*: F- CTGCAGTTGCCTCAACAAAA, R- CATGAGACTGCCAACCACAG

*Peg3*: F- TTCATCCTGAGCGAGAAGGT, R- GATGGGTTGATTTGGGTCAC

*Zdbf2*: F- AAGCGACCGACATCATCAG, R- TGGGTATGTGTGCAAATGAGA

*Mcts2*: F- GAACCAGTGGGGTAATCACG, R- CACACACAGAGATACAACGCAAT

*Phlda2*:F-TCGAAATGGCTTCGAAAATC, R- AAGAAAAGTGACATTGTTTATTTGGA

*Grb10*: F- TATGACCTCCTTGCCCACTC, R- AGAGGCCATCAAGCTTTCAA

*Tgfb1*: F- GCCCCAGAGTCTGAGACCA, R- ACGCGGGTGACCTCTTTAG

*Zfat*: F- CCCAGTTCCAACCACACAGT, R- CAGTGTCCTTGCACTGAGGA

*Trp73*: F- GCCCATCAAAGAGGAGTTCA, R- GCTCCTCTGTGCTTCTCCAC

*Actb*: F- GCGCAAGTACTCTGTGTGGA, R- CTGGCTGCCTCAACACCT

*Gapdh*: F- GTCACCAGGGCTGCCATT, R- GGCCTTCTCCATGGTGGT

*Ppib*: F- ATGTGGTACGGAAGGTGGAG, R- CCACTTTATTGATCAGCATTGG

*B2m*: F- TGAAGATTCATTTGAACCTGCTTA, R- AAAAGCAGAAGTAGCCACAGG

### DNA-FISH assay

NPCs were cultured and fixed as described for the nascent RNA-FISH assay. Cells were rinsed in 1x PBS three times (5 min each wash) and permeabilized by adding ice-cold 0.5% Triton X-100 in 1x PBS and incubating on ice for 10 minutes. Cells were washed again with 1x PBS three times (5 minutes each wash) and equilibrated in 2x SSC, changing the solution twice for 5 minutes each. Cells were then incubated in 2x SSC containing 0.1 mg/ml RNase A for 1 hour at 37°C. Hybridization probe preparation: Probe mix was prepared by combining 10 μg of mouse Cot-1 DNA, 5 μl salmon sperm DNA, 5 μl of yeast tRNA, and 40 ng of nick-translated probe in a microcentrifuge tube and adjusting the final volume to 20 μl with ddH_2_O. The mixture was gently tapped to mix and dehydrated in a SpeedVac at 65 °C for 30 minutes. Ten microliters of deionized formamide was added to the dried pellet, resuspended gently, and incubated at 37°C for 60 minutes with gentle shaking until fully dissolved. After the RNase treatment, cells were washed twice (5 minutes each) with 2x SSC at 42°C. A 70% deionized formamide in 2x SSC (pre-warmed to 80°C) was added to each well, and 24-well MatTek plates were floated on 80°C water bath and incubated for 3 minutes. The solution was removed as quickly as possible, and ice-cold (−20°C) 70% ethanol was added to each well and incubated for 5 minutes on ice. Cells were dehydrated sequentially in 95% ethanol (5 minutes), followed by 100% ethanol (5 minutes), and kept in 100% ethanol until proceeding to hybridization. Immediately before the hybridization, the probe mixture was denatured at 95°C for 10 minutes and cooled on ice for 2 minutes. Ten microliters of pre-warmed 2X hybridization buffer (20% Dextran Sulfate, 4x SSC in dH_2_O at 37°C) was added to 10 μl of probe and 20 μl of the final probe mixture was added to each well. Plates were sealed with Parafilm and incubated overnight at 37°C in a humidified chamber. After probe hybridization, the probe mixture was removed, and 50% deionized formamide in 2X SSC (pH 7.0) was added to each well and incubated at 37°C for 30 minutes. Cells were then washed sequentially with 2x SSC (15 minutes at 37°C), 1x SSC (15 minutes at 37°C), and equilibrated in 4x SSC for 5 minutes at RT. DAPI diluted 1:10,000 in 4x SSC was added to each well and incubated for 3 minutes at RT, followed by two washes with 4x SSC (5 minutes each). A total 100 μl of ibidi mounting medium (ibidi GmbH, Cat.No.50001) was added to each well, and plates were wrapped in aluminium foil and stored at 4°C until imaging.

Primers used for PCR amplification ( probes for nick translation):

Cth Nik F: GGTCAAGTGCTTGCAACAGA

Cth Nik R: GAGCTTCCGGAGACACAGTC

Ttc4 Nik F: CTCCATGGGAGGATGAGAAA

Ttc4 Nik R: CCAAACCAAACCAAACCAAC

### Chromatin Immunoprecipitation by Sun1 and Sequencing

ESCs and NPCs at ∼80% confluence were harvested by trypsinization, rinsed and resuspended in Knockout DMEM containing 10% FBS (Gibco, Cat.no.10829-018). Crosslinking was performed by incubating 5X10^7^ cells 20 ml of Knockout DMEM containing 1% Formaldehyde and 10% FBS for 10 minutes at RT while gently shaking. Crosslinking was quenched by adding 2 ml of 1.5M glycine and incubating for an additional 10 minutes at RT with gentle shaking. Cells were centrifuged at 2000 rpm for 6 min at 4°C, medium was removed, and cell plates were washed twice with 20 ml of ice-cold 1x PBS. Cell pellets were then resuspended in 500 μl of ChIP buffer (0.1% SDS, 10 mM EDTA, 50mM Tris-HCl, pH 7.5, 1 mM PMSF in dH_2_O) and kept on ice. Chromatin was sheared using a Bioruptor® Pico sonication device (diagenode, Cat.No. B01060010) with 30 seconds ON/ 30 seconds OFF for 15 cycles. After sonication, samples were centrifuged at 10,000 g for 10 minutes at 4°C. Supernatants were transferred to 1.5 ml Eppendorf microcentrifuge tubes and stored at -80°C until use. To generate Sun1-ChIP nucleic-acid-depleted extracts, nuclear extracts were treated with Benzonase® Nuclease (Millipore Sigma, Cat. No. E1014-5KU) following the manufacturer’s instructions. Sheared chromatin DNA and fragments ranged from 300-500 bp, and concentrations were measured using a Nanodrop spectrophotometer. For each ChIP reaction, 150 μl of Dynabeads™ Protein G (Invitrogen, Cat. No. 10004D) were used. Beads were washed with ice-cold 0.02% Tween-20 in 1x PBS and resuspended in 150 μl of the same buffer. Sun1 antibody (0.8 μg; Proteintech, Cat. No. 24568-I-AP) or Rabbit IgG Isotype Control (Thermo Fisher, Cat. No. 10500c) was added to washed beads, and the volume was adjusted to 200 μl with 0.02% Tween-20 in 1x PBS. Bead-antibody mixtures were incubated for 2 hours at 4°C on a tube rotator. After incubation, excess liquid was removed, and beads were gently rinsed once with RIPA buffer (50 mM Tris-HCl (pH8.0), 150 mM NaCl, 2 mM EDTA (pH8.0), 1% NP-40, 0.5% sodium deoxicholate, 0.05% SDS, and 1 Protease inhibitor tablet (cOmplete Mini, EDTA-free, Cat. No. 39802300) in 10 ml of dH_2_O). For each IP, 150 μg of DNA-containing lysates was brought to 800 μl with 1x RIPA buffer, added to the bead-antibody complexes, and incubated overnight at 4°C on a tube rotator. The next day, beads and bound proteins-DNA complexes were separated from unbound lysate using a magnetic rack. Beads were washed three times with 1000 μl of 1x TE buffer (10 mM Tris-HCl (pH8.0), 2 mM EDTA in dH_2_O). For elution, 200 μl of elution buffer (50 mM NaCl, 50 mM Tris-HCl (pH7.5), 0.1 mM PMSF, 5 mM EDTA, and 0.1% SDS in ddH_2_O) was added and incubated at 31°C for 15 minutes in a heat block with shaking at 800 rpm. For paired-end sequencing, DNAs was extracted from 100 μl of elute using the Genomic DNA Clean & Concentrator™ -25 Kit (Zymo Research, Cat.No. D4064). Libraries were prepared and sequenced on a NextSeq500 platform (PE-150, mid output) at the Next Generation Sequencing Core Facility at Cold Spring Harbor Laboratory. Nucleic acids remaining in protein fractions for for Sun1 interacting pulldown assay were removed by incubating samples with Benzonase (Millipore Sigma, Cat. No.E1014) for 20 minutes on ice prior to use.

### qPCR and Sanger sequencing

One μg of genomic DNA from Sun1-ChIP assay and from the input sample was used as template. Quantitative PCR was performed using PowerUp™ SYBR® Green Master Mix (Thermo Fisher, Cat. No. A25743) and the primers covering SNPs in the *Cth* gene (F-GAGCCTGGGAGGATATGAGA, R- AAGCTCGATCCAGGTCTTCA) and *Ttc4* gene (F – GACAGGGCGGAACTATACCA, R-genes. qPCR products were Sanger sequenced using the GenScript Biotec Sanger sequencing service.

### Solution Digestion Analysis by LC-MS/MS

Fifty microliters of elute from each sample was sent to the Taplin Biological Mass Spectrometry Facility at Harvard Medical School. In brief, 10ul (20ng/μl) of modified sequencing-grade trypsin (Promega, Madison, WI) was spiked into 300 μl PBS, and samples were incubated overnight at 37°C. Samples were acidified by adding 20 μl of 20% formic acid and desalted using STAGE tip. On the day of analysis, the samples were reconstituted in 10 µl of HPLC solvent A. A nano-scale reverse-phase HPLC capillary column was prepared by packing 2.6 µm C18 spherical silica beads into a fused silica capillary (100 µm inner diameter x ∼30 cm length) with a flame-drawn tip. After column equilibration, samples were loaded using a Famos autosampler (LC Packings, San Francisco, CA) onto the column. A gradient was formed, and peptides were eluted with increasing concentrations of solvent B (97.5% acetonitrile, 0.1% formic acid). As peptides eluted, they were subjected to electrospray ionization and then entered into an LTQ Orbitrap Velos Elite ion-trap mass spectrometer (Thermo Fisher Scientific, Waltham, MA). Peptides were detected, isolated, and fragmented to generate tandem mass spectra of specific fragment ions for each peptide. Peptide sequences (and hence protein identity) were determined by matching acquired MS/MS fragmentation patterns to protein databases using Sequest software program (Thermo Fisher Scientific, Waltham, MA). All databases include a reversed version of all the sequences, and the data was filtered to between a one and two percent peptide false discovery rate.

### Allele-specific ChIP-seq analysis and motif discovery

Allele-specific ChIP-seq analysis was performed using the MEA pipeline^4^. An in silico diploid C57BL/6J / CAST/EiJ genome was reconstructed using the SHAPEIT2 pipeline by combining the mm10 reference genome (mgp.v5.merged.snps_all.dbSNP142.vcf.gz) with the corresponding genetic variants (mgp.v5.merged.indels.dbSNP142.normed.vcf.gz). Sequencing reads (.fastq) were aligned to the reconstructed in silico genome using Bowtie2 aligner (version 2.3.4.3). The resulting BAM files were processed through the ChIP-seq pipeline embedded in SeqMonk bioinformatic web portal, and peaks were called using the MACS algorithm with a p-value threshold of 0.05 and a fragment size of 500 bp. Motif analysis was performed using HOMER (v4.11) within the Basepair web portal (Version 3.2.7).

### TA-bridge *in vivo* visualization, interacting protein pull-down, and mass spectrometry

mESCs were seeded at 0.4x10^4^ cells per well in 8-well ibidi chamber slides and incubated for 24 hours prior to transfection with biotinylated DNA oligos. For immunofluorescence assays, 0.03 μg of biotinylated oligos per well were used. For *in vivo* TA-bridge interacting protein pull-down, 2.5x10^5^ cells per well were seeded in three 6-well plates 24 hrs before transfection. A total of 0.5 μg of biotinylated oligos per well was transfected using the Xfect™ transfection reagent kit (Clonetech, Cat.No. 631317) according to the manufacturer’s instructions. Cells were incubated overnight and then fixed with 4% PFA and stored at 4°C until use for imaging. For TA-bridges interacting protein pull-down, cells were rinsed three times with 1x PBS and nuclear proteins were extracted using the nuclear extraction kit (ActiveMotif®, Cat.No. 40010). For each pull-down reaction, 200 μl of Dynabeads^TM^ MyOne Streptavidin T1 and 200 μg of nuclear extract were used. Beads were washed three times with 500 μl of 2x binding buffer and resuspended in 150 μl of 2x B&W buffer. The nuclear extract volume was adjusted to 150 μl by adding Complete Lysis Buffer (from the nuclear extract kit) and combined with the beads. Binding was carried out at RT for 1 hour on a tube rotator. After incubation, tubes were briefly centrifuged and supernatants were removed using a magnatic rack. Beads were washed four times with 1 ml of 1x B&W buffer and then resuspended in 20 μl elution buffer (50mM NaCl, 50mM Tris-HCl (pH7.5), 0.1 mM PMFS, 5 mM EDTA, 0.1% SDS in Milli-Q water) and boiled at 95°C for 7 min while gently vortexing in a heat rack. Tubes were briefly centrifuged and elutes were collected using a magnetic rack. Elutes were stored at -80°C until analysis. For MS analysis, 60 μl (20 μl combined from 3 preps) was used.

### Identification of proteins bound to TA-bridges by MS

MS was performed at the Cold Spring Harbor laboratory Mass Spectrometry Core Facility. On-membrane proteolysis was carried out as follows. Excised nylon membranes were hydrated and washed three times with 500 ul of 100 mM ammonium bicarbonate (ABC) for 10 minutes at 55°C under agitation. Next, 500 μl of fresh 3 mM TCEP (Tris(2-carboxyethyl)phosphine hydrochloride)/50mM ABC was added and incubated at 55°C for 20 minutes while vortexing at 500 rpm. The TCEP solution was removed, and 500 μl of fresh 10 mM CEMTS (2-carboxymethyl methanethiosulfonate) in 10% ethanol was added. Membranes were incubated at 55°C for 5 minutes while vortexing at 500 rpm. The CEMTS solution was removed, and membranes were washed three times with 500 μl of 50 mM ABC for 5 minutes at 55°C with agitation. Trypsin solution (100 ng/μl; Sequencing-grade modified porcine trypsin, Promega) was prepared in 50 mM ABC. Five microliters of trypsin solution was added directly onto each membrane piece, followed by 200 microliters of trypsin solution was added into the middle of the membrane piece, followed by 200 μl of 50 mM ABC containing 0.1% ProteaseMax surfactant. Membranes were digested overnight at 37°C while vortexing at 500 rpm. After digestion, 200 μl of 80% acetonitrile (CAN)/1% TFA was added and incubated for 15 minutes at 55°C with agitation. Elutes were collected into clean tubes, lyophilized by vacuum centrifugation, and the peptide pellets were resuspended in 20 μ of loading buffer (55% DMSO, 0.1% TFA in water). Peptides were loaded on a 30 cm x 75 μM inner-diameter column packed with Reprosil 1.9 μm C18 silica particles and resolved using a 5-35% acetonitrile gradient in water (0.1% formate) at a flow rate of 250 nl/minute. Eluting peptides were ionized by electrospray (2200V) and analyzed using an Exploris 480 mass spectrometer (Thermo). The instrument was operated to acquire 120K-resolution MS1 scans (m/z 380-2000 Th) and 60K HCD MS2 spectra at stepped normalized collision energies of 28,33 and 38%, with the first mass locked to 100 Th. Raw files were searched using the Mascot scoring engine within ProteomeDiscoverer, with mass tolerances set at 5 ppm for MS1 and 0.01 Da for MS2. Spectra were matched against the UniProt mouse consensus database supplemented with the common contaminants atabase (cRAP). Methionine oxidation and N/Q deamidation were set as variable modifications. Peptide spectral matches were filtered to maintain FDR < 1%.

### Electrophoretic mobility shift assay (EMSA)

EMSA was performed using LightShift® Chemiluminescent EMSA Kit (Thermo Scientific, Cat.No. 20148). Biotinylated oligos were synthesized by bioSYNTHESIS^®^. The single-strandard biotinylated oligos usd were: BM-1: TAAATAAATAAATAAATAAATAAATAAATAAA, BM-2: ATTTATTTATTTATTTATTTATTTATTTATTT, Control: GCAT. Oligos were tagged at the 3’ end with Desthiobiotin-TEG, separated from the oligo sequence by six -Sp18- spacers. Double-stranded TA-bridges were generated by anneling BM-1 with a non-biotinilated complimentary oligo, and purified using the Genomic DNA Clean & Concentrator™ (ZYMO RESEARCH, Cat. No. D4065). A total of 30 fmol of each biotinylated oligo was used per reaction. Biotinylated Epstein-Barr nuclear antigen (EBNA) DNA was used as a negative control. 6% native polyacrylamide gels (1.0 mm, 8x10 cm) were used for gel-shift assay. Nuclear extracts were prepared from 70% confluent ESC cultures using the nuclear extraction kit (ActiveMotif®, Cat.No. 40010). Binding reactions were assembled by combining 10x binding buffer, 1 μl poly(dI.dC), Nuclear extract protein, and biotinylated DNA oligos to finl volume of 20 μL. Binding mixture were incubated for 20 minutes at RT. Then, 5 μl of 5x loading buffer was added to each reaction and mixed by pipetting. Prior to sample loading, native gels were pre-run for 60 minutes at 100 V. Samples were electrophoresed at 100 V until the bromophenol blue dye migrated to approximately three-quarters of the gel length. For protein transfer, Biodyne® A pre-Cut membranes (Thermo Scientific, Cat.No. 77015) were soaked in 0.5X TBE for 10 minutes, and gels, nylon membranes, and blotting paper were assembled as specified by the manufacturer. Transfers were performed in ice-cold 0.5X TBE, with the transfer apparatus fully submerged on ice, for 45 minutes at 380 mA. Nylon membranes were semi-dried on a paper towel with the staining side facing upward. Crosslinking was performed using a UV StrataLinker™ 1800 (Stratagene) at 120,000 μJ for 60 seconds. Membranes were transferred directly to blocking buffer pre-warmed to 37°C and incubated for 15 minutes in the blocking buffer on a slow rocker. Membranes were then rinsed once in 20 ml 1x wash buffer and washed four times for 5 minutes each 20 ml 1x wash buffer with gentle shaking. Membranes were transferred to a clean container and incubated in substrate equilibration buffer for 5 minutes. Excess buffer was removed by blotting, and 12 ml of substrate working solution was added and incubated for 5 minutes without shaking. After removing excess substrate (without over-drying), membranes were wrapped in plastic film and exposed to X-ray film (Amersham Hyperfilm™ ECL, GE Healthcare Limited, Cat.No. 28906836) for 7 minuits.

## Data availability

The Sun1 ChIP-seq data (raw and processed) supporting the finding of this study have been deposited in the Gene Expression Omnibus (GEO) under accession code GSE228344. All other data supporting the finding of this study has been submitted with the manuscript.

## Acknowledgements

We wold like to thank the members of the Spector laboratory for their critical discussions and advice throughout this study. We thank Prof. Min Han for generously providing the Sun1 antibody. We thank Osama El Demerdash for assistance in establishing the computational pipeline, Tse-Luen Wee and Nour El-Amine for assistance with microscopy, and Andrew Barazia (Intelligent Imaging Innovation- 3i) for enabling use of the 3i lattice light-sheet microscope for live imaging experiments. We would also thank Paolo Cifani (CSHL) and Ross Tomaino (Taplin Mass Spectrometry Facility, Harvard Medical School) for assisting with mass spectrometry experiments, and Sara Goodwin (CSHL) for assisting with DNA sequencing. We acknowledge the CSHL Cancer Center Shared Resources (Microscopy, Mass Spectrometry, Next-Gen Sequencing) for services and technical expertise (NCI 2P3OCA45508). Sequencing was performed using equipment purchased through NIH grant S10OD028632-01. This research was supported by R35GM131833 (D.L.S.) and grant 18-26 from the Charles H. Revson Foundation (G.I.B.).

## Author Contribution

G.I.B. and D.L.S. conceived and designed the overall study. G.I.B. designed and performed all the experiments and analysis. G.I.B. and D.L.S. wrote and edited the manuscript.

## References

1 Bickmore, W. A. The spatial organization of the human genome. Annu Rev Genomics Hum Genet 14, 67–84, doi:10.1146/annurev-genom-091212-153515 (2013).

2 Binder, H. et al. Transcriptional regulation by histone modifications: towards a theory of chromatin re-organization during stem cell differentiation. Phys Biol 10, 026006, doi:10.1088/1478-3975/10/2/026006 (2013).

3 Cremer, T. & Cremer, M. Chromosome Territories. Csh Perspect Biol 2, doi:ARTN a003889 10.1101/cshperspect.a003889 (2010).

4 Fraser, P. & Bickmore, W. Nuclear organization of the genome and the potential for gene regulation. Nature 447, 413–417, doi:10.1038/nature05916 (2007).

5 Hubner, M. R., Eckersley-Maslin, M. A. & Spector, D. L. Chromatin organization and transcriptional regulation. Curr Opin Genet Dev 23, 89–95, doi:10.1016/j.gde.2012.11.006 (2013).

6 Meaburn, K. J. & Misteli, T. Cell biology: chromosome territories. Nature 445, 379–781, doi:10.1038/445379a (2007).

7 Takizawa, T., Meaburn, K. J. & Misteli, T. The meaning of gene positioning. Cell 135, 9–13, doi:10.1016/j.cell.2008.09.026 (2008).

8 Abed, J. A. et al. Highly structured homolog pairing reflects functional organization of the Drosophila genome. Nat Commun 10, doi:ARTN 4485 10.1038/s41467-019-12208-3 (2019).

9 Duncan, I. W. Transvection effects in Drosophila. Annu Rev Genet 36, 521–556, doi:10.1146/annurev.genet.36.060402.100441 (2002).

10 Galouzis, C. C. & Prud’homme, B. Transvection regulates the sex-biased expression of a fly X-linked gene. Science 371, 396-+, doi:10.1126/science.abc2745 (2021).

11 Hogan, M. S., Parfitt, D. E., Zepeda-Mendoza, C. J., Shen, M. M. & Spector, D. L. Transient Pairing of Homologous Oct4 Alleles Accompanies the Onset of Embryonic Stem Cell Differentiation. Cell Stem Cell 16, 275–288, doi:10.1016/j.stem.2015.02.001 (2015).

12 Jack, J. W. & Judd, B. H. Allelic pairing and gene regulation: A model for the zeste-white interaction in Drosophila melanogaster. Proc Natl Acad Sci U S A 76, 1368–1372, doi:10.1073/pnas.76.3.1368 (1979).

13 Viets, K. et al. Characterization of Button Loci that Promote Homologous Chromosome Pairing and Cell-Type-Specific Interchromosomal Gene Regulation. Dev Cell 51, 341-+, doi:10.1016/j.devcel.2019.09.007 (2019).

14 D’Angelo, M. A. Nuclear pore complexes as hubs for gene regulation. Nucleus-Phila 9, 142–148, doi:10.1080/19491034.2017.1395542 (2018).

15 Mao, Y. T. S., Zhang, B. & Spector, D. L. Biogenesis and function of nuclear bodies. Trends Genet 27, 295–306, doi:10.1016/j.tig.2011.05.006 (2011).

16 Meldi, L. & Brickner, J. H. Compartmentalization of the nucleus. Trends Cell Biol 21, 701–708, doi:10.1016/j.tcb.2011.08.001 (2011).

17 Quinodoz, S. A. et al. Higher-Order Inter-chromosomal Hubs Shape 3D Genome Organization in the Nucleus. Cell 174, 744-+, doi:10.1016/j.cell.2018.05.024 (2018).

18 Zimber, A., Nguyen, Q. D. & Gespach, C. Nuclear bodies and compartments: Functional roles and cellular signalling in health and disease. Cell Signal 16, 1085–1104, doi:10.1016/j.cellsig.2004.03.020 (2004).

19 Eckersiey-Maslin, M. A. & Spector, D. L. Random monoallelic expression: regulating gene expression one allele at a time. Trends Genet 30, 237–244, doi:10.1016/j.tig.2014.03.003 (2014).

20 Ferguson-Smith, A. C. Genomic imprinting: the emergence of an epigenetic paradigm. Nat Rev Genet 12, 565–575, doi:10.1038/nrg3032 (2011).

21 Reik, W. & Walter, J. Imprinting mechanisms in mammals. Curr Opin Genet Dev 8, 154–164, doi:10.1016/s0959-437x(98)80136-6 (1998).

22 Balasooriya, G. I. & Spector, D. L. Allele-specific differential regulation of monoallelically expressed autosomal genes in the cardiac lineage. Nat Commun 13, 5984, doi:10.1038/s41467-022-33722-x (2022).

23 Chen, M. et al. Live imaging of RNA and RNA splicing in mammalian cells via the dcas13a-SunTag-BiFC system. Biosens Bioelectron 204, 114074, doi:10.1016/j.bios.2022.114074 (2022).

24 Daigle, N. & Ellenberg, J. LambdaN-GFP: an RNA reporter system for live-cell imaging. Nat Methods 4, 633–636, doi:10.1038/nmeth1065 (2007).

25 Haim, L., Zipor, G., Aronov, S. & Gerst, J. E. A genomic integration method to visualize localization of endogenous mRNAs in living yeast. Nat Methods 4, 409–412, doi:10.1038/nmeth1040 (2007).

26 Hocine, S., Raymond, P., Zenklusen, D., Chao, J. A. & Singer, R. H. Single-molecule analysis of gene expression using two-color RNA labeling in live yeast. Nat Methods 10, 119–121, doi:10.1038/nmeth.2305 (2013).

27 Janicki, S. M. et al. From silencing to gene expression: real-time analysis in single cells.Cell 116, 683–698, doi:10.1016/s0092-8674(04)00171-0 (2004).

28 Ma, H. H. et al. Multiplexed labeling of genomic loci with dCas9 and engineered sgRNAs using CRISPRainbow. Nature Biotechnology 34, 528–530, doi:10.1038/nbt.3526 (2016).

29 Maass, P. G. et al. Spatiotemporal allele organization by allele-specific CRISPR live-cell imaging (SNP-CLING). Nat Struct Mol Biol 25, 176-+, doi:10.1038/s41594-017-0015-3 (2018).

30 Tsukamoto, T. et al. Visualization of gene activity in living cells. Nat Cell Biol 2, 871–878, doi:10.1038/35046510 (2000).

31 Vera, M., Biswas, J., Senecal, A., Singer, R. H. & Park, H. Y. Single-Cell and Single-Molecule Analysis of Gene Expression Regulation. Annual Review of Genetics, Vol 50 50, 267–291, doi:10.1146/annurev-genet-120215-034854 (2016).

32 Chen, H. M. & Larson, D. R. What have single-molecule studies taught us about gene expression? Gene Dev 30, 1796–1810, doi:10.1101/gad.281725.116 (2016).

33 Shao, S. P., Xue, B. X. & Sun, Y. J. Intranucleus Single-Molecule Imaging in Living Cells. Biophys J 115, 181–189, doi:10.1016/j.bpj.2018.05.017 (2018).

34 Shav-Tal, Y., Singer, R. H. & Darzacq, X. Imaging gene expression in single living cells. Nat Rev Mol Cell Bio 5, 855–862, doi:10.1038/nrm1494 (2004).

35 Blake, W. J., Kaern, M., Cantor, C. R. & Collins, J. J. Noise in eukaryotic gene expression. Nature 422, 633–637, doi:10.1038/nature01546 (2003).

36 Larson, D. R., Singer, R. H. & Zenklusen, D. A single molecule view of gene expression. Trends Cell Biol 19, 630–637, doi:10.1016/j.tcb.2009.08.008 (2009).

37 Wang, L., Frei, M. S., Salim, A. & Johnsson, K. Small-Molecule Fluorescent Probes for Live-Cell Super-Resolution Microscopy. J Am Chem Soc 141, 2770–2781, doi:10.1021/jacs.8b11134 (2019).

38 Joyce, E. F., Erceg, J. & Wu, C. T. Pairing and anti-pairing: a balancing act in the diploid genome. Curr Opin Genet Dev 37, 119–128, doi:10.1016/j.gde.2016.03.002 (2016).

39 Blattner, G., Cavazza, A., Thrasher, A. J. & Turchiano, G. Gene Editing and Genotoxicity: Targeting the Off-Targets. Front Genome Ed 2, 613252, doi:10.3389/fgeed.2020.613252 (2020).

40 Bohne, J. & Cathomen, T. Genotoxicity in gene therapy: An account of vector integration and designer nucleases. Curr Opin Mol Ther 10, 214–223 (2008).

41 Yoon, K. H. et al. Olfactory receptor genes expressed in distinct lineages are sequestered in different nuclear compartments. P Natl Acad Sci USA 112, E2403–E2409, doi:10.1073/pnas.1506058112 (2015).

42 Armelin-Correa, L. M., Gutiyama, L. M., Brandt, D. Y. C. & Malnic, B. Nuclear compartmentalization of odorant receptor genes. P Natl Acad Sci USA 111, 2782–2787, doi:10.1073/pnas.1317036111 (2014).

43 Clowney, E. J. et al. Nuclear Aggregation of Olfactory Receptor Genes Governs Their Monogenic Expression. Cell 151, 724–737, doi:10.1016/j.cell.2012.09.043 (2012).

44 Gundersen, G. G. & Worman, H. J. Nuclear Positioning. Cell 152, 1376–1389, doi:10.1016/j.cell.2013.02.031 (2013).

45 Haque, F. et al. SUN1 interacts with nuclear lamin A and cytoplasmic nesprins to provide a physical connection between the nuclear lamina and the cytoskeleton. Mol Cell Biol 26, 3738–3751, doi:10.1128/Mcb.26.10.3738-3751.2006 (2006).

46 Liu, Q. et al. Functional association of Sun1 with nuclear pore complexes. J Cell Biol 178, 785–798, doi:10.1083/jcb.200704108 (2007).

47 Li, P. et al. The function of the inner nuclear envelope protein SUN1 in mRNA export is regulated by phosphorylation. Sci Rep-Uk 7, doi:ARTN 9157 10.1038/s41598-017-08837-7 (2017).

48 Summer, M. C. & Brickner, J. The Nuclear Pore Complex as a Transcription Regulator. Csh Perspect Biol 14, doi:ARTN a039438 10.1101/cshperspect.a039438 (2022).

49 Matsumoto, A. et al. Loss of the integral nuclear envelope protein SUN1 induces alteration of nucleoli. Nucleus-Phila 7, 68–83, doi:10.1080/19491034.2016.1149664 (2016).

50 van Steensel, B. & Belmont, A. S. Lamina-Associated Domains: Links with Chromosome Architecture, Heterochromatin, and Gene Repression. Cell 169, 780–791, doi:10.1016/j.cell.2017.04.022 (2017).

51 Gonzalez-Sandoval, A. & Gasser, S. M. On TADs and LADs: Spatial Control Over Gene Expression. Trends Genet 32, 485–495, doi:10.1016/j.tig.2016.05.004 (2016).

52 Guelen, L. et al. Domain organization of human chromosomes revealed by mapping of nuclear lamina interactions. Nature 453, 948–951, doi:10.1038/nature06947 (2008).

53 Peric-Hupkes, D. et al. Molecular maps of the reorganization of genome-nuclear lamina interactions during differentiation. Mol Cell 38, 603–613, doi:10.1016/j.molcel.2010.03.016 (2010).

54 Leemans, C. et al. Promoter-Intrinsic and Local Chromatin Features Determine Gene Repression in LADs. Cell 177, 852–864 e814, doi:10.1016/j.cell.2019.03.009 (2019).

55 Luo, L. et al. The nuclear periphery of embryonic stem cells is a transcriptionally permissive and repressive compartment. J Cell Sci 122, 3729–3737, doi:10.1242/jcs.052555 (2009).

56 Wu, F. & Yao, J. Identifying Novel Transcriptional and Epigenetic Features of Nuclear Lamina-associated Genes. Sci Rep 7, 100, doi:10.1038/s41598-017-00176-x (2017).

57 Bonev, B. et al. Multiscale 3D Genome Rewiring during Mouse Neural Development. Cell 171, 557-+, doi:10.1016/j.cell.2017.09.043 (2017).

58 Monahan, K., Horta, A. & Lomvardas, S. LHX2-and LDB1-mediated trans interactions regulate olfactory receptor choice. Nature 565, 448-+, doi:10.1038/s41586-018-0845-0 (2019).

59 Dekker, J. et al. The 4D nucleome project. Nature 549, 219–226, doi:10.1038/nature23884 (2017).

60 Reiff, S. B. et al. The 4D Nucleome Data Portal as a resource for searching and visualizing curated nucleomics data. Nat Commun 13, doi:ARTN 2365 10.1038/s41467-022-29697-4 (2022).

61 Shevelyov, Y. Y. & Ulianov, S. V. The Nuclear Lamina as an Organizer of Chromosome Architecture. Cells 8, doi:ARTN 136 10.3390/cells8020136 (2019).

62 Czapiewski, R., Robson, M. I. & Schirmer, E. C. Anchoring a Leviathan: How the Nuclear Membrane Tethers the Genome. Front Genet 7, doi:ARTN 82 10.3389/fgene.2016.00082 (2016).

63 Snyder, M. J. et al. Anchoring of Heterochromatin to the Nuclear Lamina Reinforces Dosage Compensation-Mediated Gene Repression. Plos Genet 12, doi:ARTN e1006341 10.1371/journal.pgen.1006341 (2016).

64 Rivera-Mulia, J. C. et al. Allele-specific control of replication timing and genome organization during development. Genome Research 28, 800–811, doi:10.1101/gr.232561.117 (2018).

65 Arao, Y. et al. A+U-rich-element RNA-binding factor 1/heterogeneous nuclear ribonucleoprotein D gene expression is regulated by oestrogen in the rat uterus. Biochem J 361, 125–132, doi:Doi 10.1042/0264-6021:3610125 (2002).

66 Wu, F. N. & Yao, J. Identifying Novel Transcriptional and Epigenetic Features of Nuclear Lamina-associated Genes. Sci Rep-Uk 7, doi:ARTN 100 10.1038/s41598-017-00176-x (2017).

67 Leemans, C. et al. Promoter-Intrinsic and Local Chromatin Features Determine Gene Repression in LADs. Cell 177, 852-+, doi:10.1016/j.cell.2019.03.009 (2019).

68 Wu, C. H. et al. NELF and DSIF cause promoter proximal pausing on the hsp70 promoter in Drosophila. Gene Dev 17, 1402–1414, doi:10.1101/gad.1091403 (2003).

69 Gressel, S. et al. CDK9-dependent RNA polymerase II pausing controls transcription initiation. Elife 6, doi:ARTN e29736 10.7554/eLife.29736 (2017).

70 Nazer, E., Dale, R. K., Chinen, M., Radmanesh, B. & Lei, E. P. Argonaute2 and LaminB modulate gene expression by controlling chromatin topology. Plos Genet 14, doi:ARTN e1007276 10.1371/journal.pgen.1007276 (2018).

71 Hah, J. & Kim, D. H. Deciphering Nuclear Mechanobiology in Laminopathy. Cells 8, doi:10.3390/cells8030231 (2019).

72 Chen, Y. et al. The SUN1-SPDYA interaction plays an essential role in meiosis prophase I. Nat Commun 12, 3176, doi:10.1038/s41467-021-23550-w (2021).

73 Hoyt, S. J. et al. From telomere to telomere: The transcriptional and epigenetic state of human repeat elements. Science 376, eabk3112, doi:10.1126/science.abk3112 (2022).

74 Wang, Y., Zhao, Y., Bollas, A., Wang, Y. & Au, K. F. Nanopore sequencing technology, bioinformatics and applications. Nat Biotechnol 39, 1348–1365, doi:10.1038/s41587-021-01108-x (2021).

75 Bzymek, M. & Lovett, S. T. Instability of repetitive DNA sequences: the role of replication in multiple mechanisms. Proc Natl Acad Sci U S A 98, 8319–8325, doi:10.1073/pnas.111008398 (2001).

76 Altemose, N. et al. DiMeLo-seq: a long-read, single-molecule method for mapping protein-DNA interactions genome wide. Nat Methods 19, 711–723, doi:10.1038/s41592-022-01475-6 (2022).

77 Nguyen, G. H. et al. Regulation of gene expression by the BLM helicase correlates with the presence of G-quadruplex DNA motifs. Proc Natl Acad Sci U S A 111, 9905–9910, doi:10.1073/pnas.1404807111 (2014).

78 Mao, S. Q. et al. DNA G-quadruplex structures mold the DNA methylome. Nat Struct Mol Biol 25, 951–957, doi:10.1038/s41594-018-0131-8 (2018).

79 Tye, B. K. MCM proteins in DNA replication. Annu Rev Biochem 68, 649–686, doi:10.1146/annurev.biochem.68.1.649 (1999).

80 Roca, J. Topoisomerase II: a fitted mechanism for the chromatin landscape. Nucleic Acids Res 37, 721–730, doi:10.1093/nar/gkn994 (2009).

81 Dixon, J. R. et al. Chromatin architecture reorganization during stem cell differentiation. Nature 518, 331–336, doi:10.1038/nature14222 (2015).

82 Winick-Ng, W. et al. Cell-type specialization is encoded by specific chromatin topologies. Nature 599, 684-+, doi:10.1038/s41586-021-04081-2 (2021).

83 Chess, A., Simon, I., Cedar, H. & Axel, R. Allelic inactivation regulates olfactory receptor gene expression. Cell 78, 823–834, doi:10.1016/s0092-8674(94)90562-2 (1994).

84 Singh, N. et al. Coordination of the random asynchronous replication of autosomal loci. Nat Genet 33, 339–341, doi:10.1038/ng1102 (2003).

85 Lionnet, T. et al. A transgenic mouse for in vivo detection of endogenous labeled mRNA. Nat Methods 8, 165–170, doi:10.1038/nmeth.1551 (2011).

